# An Agent-Based Model of Metabolic Signaling Oscillations in *Bacillus subtilis* Biofilms

**DOI:** 10.1101/2024.12.20.629727

**Authors:** Obadiah J. Mulder, Maya Peters Kostman, Abdulrahmen Almodaimegh, Michael D. Edge, Joseph W. Larkin

## Abstract

Microbes of nearly every species can form biofilms, communities of cells bound together by a self-produced matrix. It is not understood how variation at the cellular level impacts putatively beneficial, colony-level behaviors, such as cell-to-cell signaling. Here we investigate this problem with an agent-based computational model of metabolically driven electrochemical signaling in *Bacillus subtilis* biofilms. In this process, glutamate-starved interior cells release potassium, triggering a depolarizing wave that spreads to exterior cells and limits their glutamate uptake. More nutrients diffuse to the interior, temporarily reducing glutamate stress and leading to oscillations. In our model, each cell has a membrane potential coupled to metabolism. As a simulated biofilm grows, collective membrane potential oscillations arise spontaneously as cells deplete nutrients and trigger potassium release, reproducing experimental observations. We further validate our model by comparing spatial signaling patterns and cellular signaling rates with those observed experimentally. By oscillating external glutamate and potassium, we find that biofilms synchronize to external potassium more strongly than to glutamate, providing a potential mechanism for previously observed biofilm synchronization. By tracking cellular glutamate concentrations, we find that oscillations evenly distribute nutrients in space: non-oscillating biofilms have an external layer of well-fed cells surrounding a starved core, whereas oscillating biofilms exhibit a relatively uniform distribution of glutamate. Our work shows the potential of agent-based models to connect cellular properties to collective phenomena and facilitates studies of how inheritance of cellular traits can affect the evolution of group behaviors.

## Introduction

Bacterial biofilms are large communities of cells that exist in nearly every environment [10]. They are bound together by an extracellular matrix that provides both stability and protection [3, 8, 1]. Biofilms exhibit a variety of emergent behaviors that give biofilm-dwelling microbes advantages unavailable to planktonic cells [16, 31, 42]. For example, cells within biofilms differentiate into heterogeneous phenotypes [22, 45, 46, 21], divide labor [28, 36], and coordinate behavior via chemical signals [13, 33, 49, 23]. These group phenomena have led researchers to assert that biofilms represent a transition between single-celled and multicellular life [38, 7].

A striking multicellular behavior is the presence of cell-to-cell electrochemical signals that influence metabolism in *Bacillus subtilis* biofilms [25, 34]. As a biofilm expands, fewer nutrients penetrate to the center; most are consumed by exterior cells [43, 51]. The paucity of nutrients in the interior raises a problem: if interior cells are starved, the integrity of the biofilm is at risk [25]. *In vitro B. subtilis* biofilms exhibit a behavior that seems to allow them to navigate this challenge. When interior cells become starved, they release potassium, depolarizing nearby cells and hampering their ability to absorb glutamate. In turn, nearby cells become distressed, release potassium, and hyperpolarize, eventually leading to a wave of potassium release. This wave propagates to the biofilm exterior [34]. It has been hypothesized that glutamate consumption among cells in the exterior slows down enough that glutamate can diffuse to the center [25]. Once interior cells have enough glutamate, they cease releasing potassium, allowing exterior cells to repolarize and resume consumption, eventually leading to stress and another wave of potassium release.

These repeated waves of potassium release have been referred to as a form of microbial “signaling” [34, 29, 11]. Potassium signaling has been proposed to allocate nutrients efficiently at the colony level [25, 24], and it is heterogeneous at the cellular level. Some cells participate in signaling and hyperpolarize during signaling waves, whereas others do not [20]. It is unknown how cellular variation in signaling behavior affects biofilm-level properties, such as distributions of nutrients. In order to answer this question, we need models that can connect cell-level properties, such as signaling state and inheritance of signaling behavior, to colony-level phenomena.

Several computational models of *B. subtilis* signaling behavior have been introduced to explore hypotheses about the causes and effects of signaling. Zhai and colleagues (2019) proposed an agent-based model to explain observations they had made about signaling. Their *in vitro* experiments revealed that a roughly constant proportion of cells signal each oscillation, that the same cells tend to release potassium in repeated signaling waves, and that signaling behavior is weakly heritable—that is, daughter cells of signaling cells are more likely than average to participate in signaling waves. They modeled signaling as a percolation process in which a cell only signals during a depolarization wave if it both has a binary trait that predisposes it to signaling and is adjacent to another signaling cell in the biofilm. Using an agent-based model in which agents represent individual cells allowed them to test whether signaling in this manner would transmit a signal across the biofilm consistently. However, their model focused on small sub-regions of the biofilm to match the limitations of their experimental system—a roughly 35x230 rectangle of cells at the edge of the biofilm, where the colony is close to two-dimensional. Their model also did not include nutrient diffusion or uptake, preventing its use for studying how individual cell behaviors affect the distribution of nutrients or growth of the biofilm.

Other models of *B. subtilis* depolarization waves are based on systems of differential equations. Martinez-Corral et al. (2018) produced a model of a one-dimensional slice of the biofilm, extending from the center to an edge. Ford et al. (2021) extended this to two dimensions, simulating a complete biofilm. Both models aimed to capture signaling and nutrient patterns at the scale of an entire biofilm. These models explicitly simulate nutrient diffusion and metabolism and have signaling operate through mechanisms that depend on internal glutamate concentration, providing powerful and accurate recreation of biofilm-wide signaling dynamics. However, modeling these complex interactions at a larger scale using differential equations comes at the cost of resolution. These models describe phenomena on the scale of the biofilm but do not distinguish individual cells. Their advantages are thus opposite those of the agent-based model of Zhai and colleagues, but neither can describe the effects of individual cell behaviors on broad patterns of nutrient distribution or signaling.

The model we propose strikes a compromise between the flexibility and resolution of the agent-based approach of Zhai and colleagues (2019) and the scalability of ODE models. Our approach is agent-based, but the agent-based elements are overlaid on a simplified version of the ODE model developed by Martinez-Corral et al. (2019). Via this hybrid strategy, our model retains some of the benefits of both previous approaches. Our model enables simulation at the scale of an entire flow-cell biofilm [15, 35], comprising approximately 51,000 individual cells, each with unique potassium, glutamate, membrane potential, and signaling dynamics.

We validate our model by comparing the behaviors of simulated biofilms with those observed in experiments, including signaling patterns at local and colony-wide scales, response to various stressors, and growth patterns. We show that many of the distinctive features of *B. subtilis* signaling, including waves of depolarization and the fraction, identity, and descent of cells that participate in signaling, can emerge naturally from our model. We then demonstrate the application of our model by exploring open questions regarding synchronization of oscillations among neighboring biofilms [24] and the effect of signaling on glutamate distribution.

## Results

### Model overview

Our model aims to describe an oscillating hyperpolarization-depolarization behavior observed in *B. subtilis* biofilms grown in flow cells [25, 34]. In such scenarios, when a biofilm grows past a certain size, metabolically stressed interior cells release potassium ions. The primary source of nitrogen in flow-cell experiments is glutamate [25], and cells absorb glutamate via a transporter whose activity depends on membrane potential. This transporter is more efficient when the cell is hyperpolarized—that is, when there is a greater charge differential between the interior of the cell and the extracellular media [44]. By releasing charged potassium ions, stressed cells increase their membrane polarization and therefore their ability to absorb nutrients.

Releasing potassium ions has an additional effect of depolarizing surrounding cells. Prindle et al. (2015) hypothesized that when interior cells are extremely stressed and release a sufficiently large amount of potassium, they can depolarize surrounding cells enough to slow their nutrient uptake. If enough cells release potassium, a chain reaction can be triggered in which nearby cells become depolarized, undergo metabolic stress, and then release ions and hyperpolarize in response. Ion release can be thought of as a form of signaling, albeit one that has direct effects on cell physiology. If enough cells signal, it can lead neighboring cells to signal, causing a wave of depolarization to spread across the biofilm.

As the wave of depolarization crosses the biofilm, nutrient absorption across the entire colony slows. This eventually allows nutrients to diffuse to the interior of the biofilm and thereby reduce metabolic stress. A side effect of reduced nutrient uptake is a corresponding reduction in growth [2], particularly in exterior cells where most biofilm expansion occurs [25, 47]. After the wave of depolarization reaches the exterior of the biofilm and nutrients diffuse through the biofilm and reduce stress in the interior, growth can resume. Consequently, whereas there is consistently rapid growth when the biofilm is small, once it surpasses a threshold size—determined by nutrient concentration in the media and the biofilm’s shape and density—it transitions to periodic growth, with growth pausing when the exterior of the biofilm is depolarized.

Our model describes the signaling waves that appear to drive these oscillations in growth (Figure 1A). We developed an agent-based model that explicitly simulates each cell spatially on a two-dimensional plane. Our model is hexagonal (to mimic the approximate 6-neighbor structure of a 2D biofilm [20]), and can be run at the scale of an entire flow-cell biofilm, with a radius of approximately 145 cells. We model glutamate as diffusing into the biofilm from outside and being consumed by cells; uptake of glutamate causes a cell’s internal glutamate level to increase. When cells are below an individual-specific threshold level of internal glutamate, they release potassium ions, allowing faster glutamate uptake.

**Figure 1:**
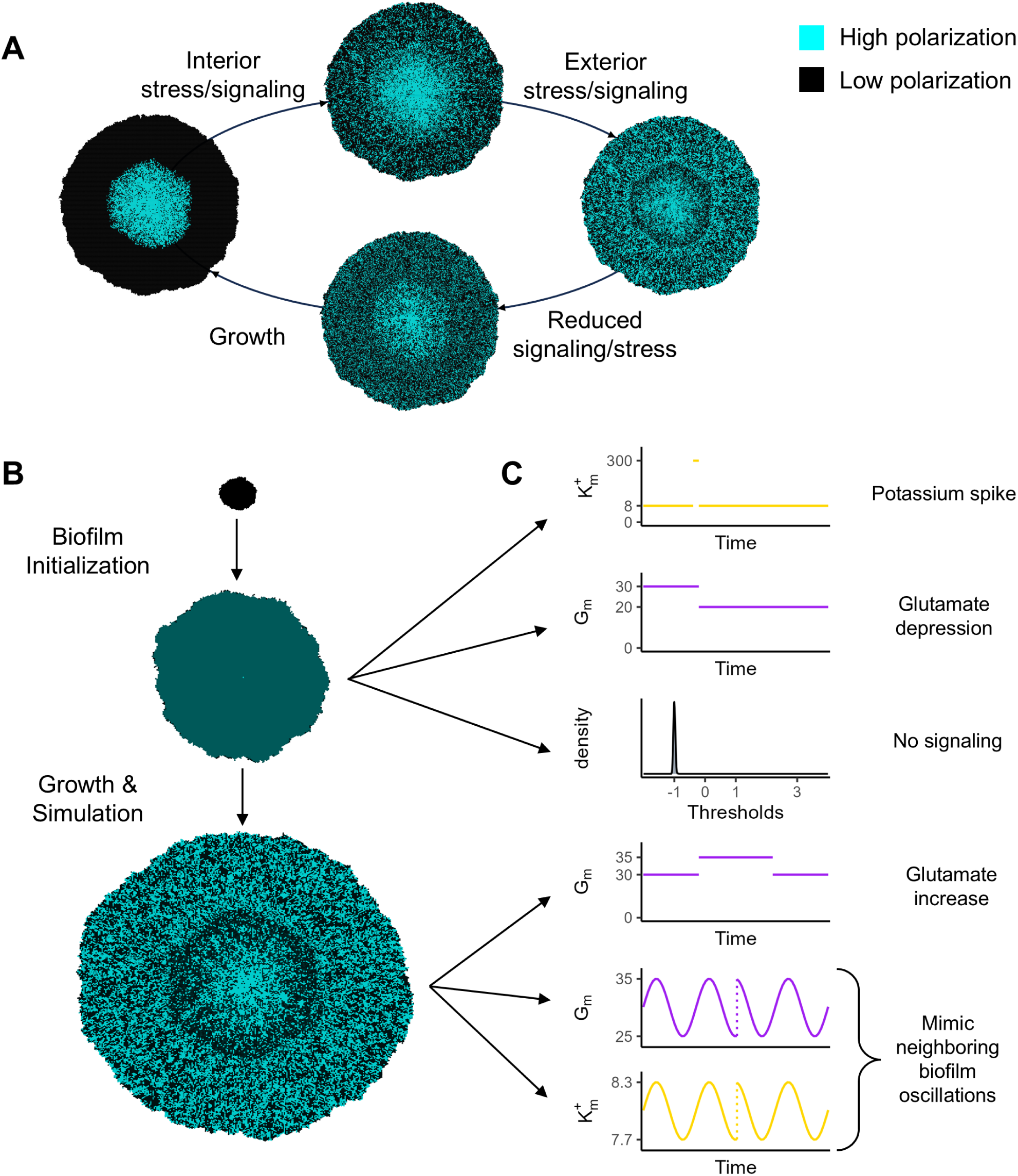
A schematic of our model. (A) shows the cycle of oscillations: growth causing interior stress, leading to signaling (indicated by cyan cells) and exterior stress, causing slowed growth and a reduction in stress, and finally back to resumed growth. (B) shows our simulation process, beginning with a very small cluster of cells, growing it for a period of time without simulating nutrients, and then growing to full size and running for many iterations with nutrient and signaling simulation. (C) shows the questions we pursue, including testing the effects of varying levels of glutamate and potassium, and of suppressing signaling.

Intracellular glutamate (*G_i_*) and potassium (*K_e_*), extracellular glutamate (*G_e_*) and potassium (*K_e_*), and cell membrane potential (*V*) are regulated by four equations taken from Martinez-Corral et al. (2019) with simplifications (equations S1, S3-S5, see Table S2 for parameter values). Each cell has a signaling threshold, *T_i_* —when a cell’s internal glutamate drops below *T_i_*, the cell signals. *T_i_* is treated as a property of the cell that remains fixed throughout the cell’s lifespan. Cells pass their signaling threshold to their offspring, with a certain amount of noise, causing signaling behavior to be partially heritable, as observed *in vitro* [50]. (We use the term “heritable” to refer to the correlation between mother and daughter cells, without assuming that the source of variation between lineages is genetic, which is unlikely in clonal biofilms.) An illustration of the potassium, glutamate, and membrane potential for a single cell during a signaling wave is shown in Figure S1.

Although the equations governing biofilm behavior are modeled on those of Martinez-Corral et al. (2019), we made modifications for use in an agent-based model. For computational tractability, we discretized coarsely with respect to time, applying the equations every time step (“tick”). A tick represents a period of approximately one minute, an interval with respect to which potassium diffuses rapidly (Supplement S3.4.1, [30]). This coarse time grid allowed us to model potassium diffusion simply by averaging it across the biofilm each tick. Glutamate diffuses more slowly than potassium [37, 34], so we model its diffusion, albeit in a simplified way (described in Supplement S3.1). Each tick, basal glutamate in the media (*G_m_*) diffuses into the biofilm and is absorbed by cells according to equation S1.

We initialized biofilms with a small number of cells such that glutamate diffused to the center easily. We then allowed them to grow to a radius of approximately 145 cells (a population of ∼51,000), at which point we stopped growth (Figure 1B). At each tick during the growth phase, we selected one-fortieth of the cells on the perimeter of the biofilm network, uniformly at random and with replacement, to reproduce. This produced growth consistent with the doubling time of *B. subtilis* (between 45 minutes and 6 hours [4, 9, 18]). Each daughter cell was placed in one of the empty nodes adjacent to the parent. Its signaling threshold was drawn from a truncated normal distribution with *—* equal to the parent’s threshold, *ff* (corresponding to the standard deviation on a non-truncated normal distribution) of 1, and bounds of [0, 3] (further described in Supplement S2). Once growth stopped, we continued the simulation for a total of 3000 ticks. This time period corresponds to approximately 48 hours, longer than *in vitro* biofilms have been observed to maintain oscillatory behavior. To replicate previous studies and make new predictions, we simulated biofilms under a variety of conditions, including reduced and increased basal glutamate, oscillated basal glutamate and potassium, and a short flood of potassium to depolarize the biofilm (Figure 1C).

### Model validation

#### Patterns of signaling

We initially explored our model by replicating behaviors and findings from previous work. We first examined whether our model produced simulated biofilms in which signaling oscillations behave similarly to *in vitro* observations. At a gross level, videos of oscillations in *in vitro* biofilms and in our simulated biofilms and reveal many similar features (videos available as files S5.1 and S5.2).

Martinez-Corral et al. (2018) observed that oscillations generally begin at a radius of 200-350 *—*m under environmental conditions identical to those in our model (30 mM glutamate). We found oscillations to start at a radius of around 110 cells (Figure 2), which corresponds to approximately 220-330 *—*m [39, 48].

**Figure 2:**
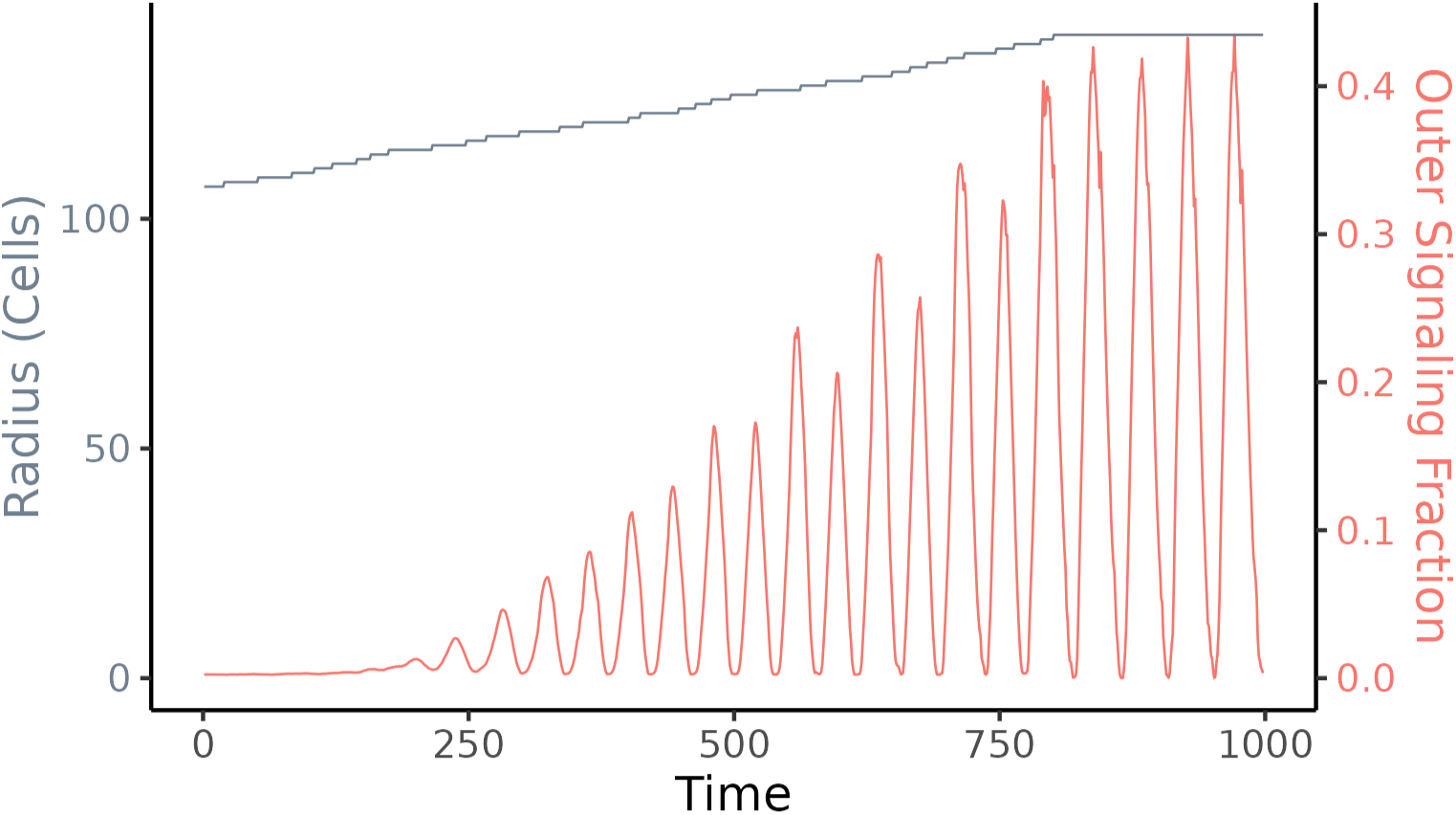
Radius (gray) and fraction of signaling cells (red) in the outer region of a simulated growing biofilm over time. The radius indicates the distance of the cell farthest from the center. Growth is limited to a radius of approximately 145 cells. A version with growth to a much larger size is shown in Figure S2, demonstrating the collapse of oscillations when the biofilm grows too large.

Zhai et al. (2019) found that near the edge of the biofilm, approximately 43% of cells are signaling during the peak of each oscillation. This observation motivated their investigation of signaling in terms of percolation theory—43% is near the minimum fraction of signaling cells that guarantees a signal moving between adjacent cells can cross the biofilm, given their other assumptions. Our simulated biofilms behave similarly, with approximately 43% of cells in the outer layers of the biofilm signaling at the height of each signaling oscillation (Figure 2).

In experimental time-lapse images of biofilm signaling, the interior and exterior of the biofilm oscillate approximately in antiphase, with the interior exhibiting much higher polarization (Supplement S5.1, Figure 3A). *In vitro*, the division between the interior and exterior (defined by oscillation) appears sharp (Figure 3B). In our simulations, we observed the same boundary (Supplement S5.2 and Figures 3D and S3). The difference in polarization can also be observed by comparing the vertical axes for inner and outer cells in Figure 3, panels A and C.

**Figure 3:**
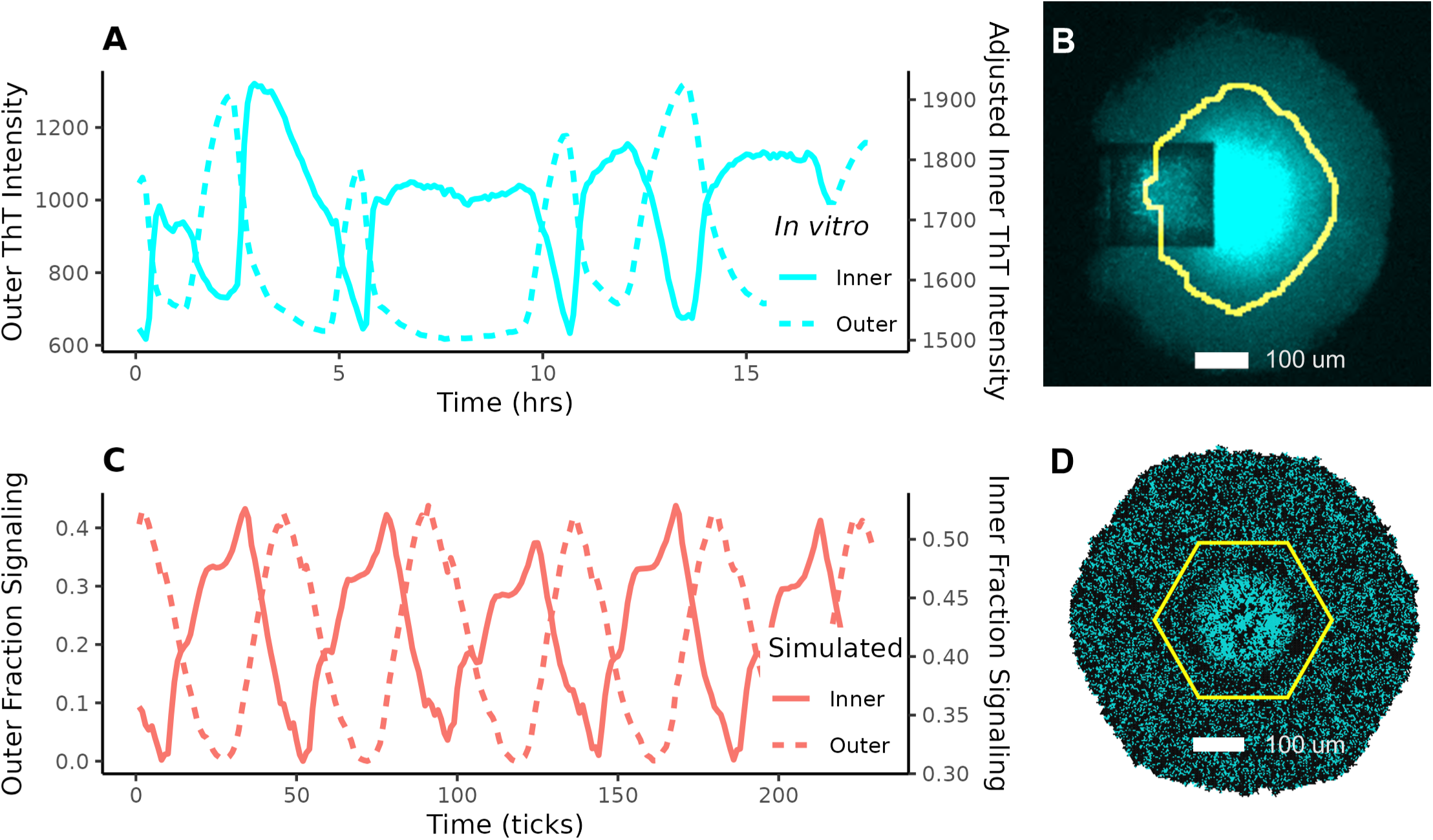
Comparison between in vitro observations of oscillations in the interior and exterior of the biofilm (A, B), and our simulations of the same (C, D). In the in vitro observations, time is given in hours and the y-axis shows the average Thioflavin-T (ThT) intensity in each region. ThT is a stain used to detect membrane polarization; polarized cells absorb it and exhibit fluorescence [19, 34]. Note that the interior has much higher ThT intensity than the exterior. (B) is an in vitro fluorescence image of a signaling biofilm (cyan represents ThT intensity; the square is a cell loading trap) and (D) is snapshot from our model, both with the boundary between inner and outer cells highlighted (yellow). In (C) and (D) cyan represents cell membrane polarization.

#### Single-cell signaling behavior

Larkin et al. (2018) found a bimodal distribution of cell-level membrane potentials during signaling peaks. Cells that had recently signaled had substantially more negative membrane potentials than those that had not. The membrane potential distribution during signaling peaks was also bimodal in our simulations, and we used the bimodality to define signaling vs. non-signaling cells, classifying those on the more highly polarized mode as signaling (Figure S4).

At the individual-cell level, signaling behavior is consistent across oscillations: cells that signal in a given wave are more likely to participate in other waves of signaling. To characterize this consistency, we used *in vitro* lineage tracing across two oscillations, again focusing only on exterior cells. We found that across a pair of oscillations, 33% of cells signaled in both waves (compared with ∼18% expected if signaling participation is independent between waves), 47% did not participate in either signaling wave, and 20% switched their signaling behavior between waves (with roughly half going either direction). These proportions are inconsistent with the null hypothesis that cell-level signaling behavior is independent between waves (Fisher’s exact test *p <* 10^−24^). We then measured pairwise consistency in our simulations to compare with our *in vitro* findings. In our simulations, we observed similar behavior, with 37% consistently signaling, 52% consistently not signaling, and 11% switching (Table 1).

**Table 1:** A comparison between individual-cell behaviors observed in vitro and those predicted by our simulations. All simulated values are for exterior cells only. Signaling fraction and recurrence rates are from Zhai et al. (2019). Signaling fraction is the maximum proportion of cells simultaneously signaling during each oscillation. The recurrence rates are the probabilities that a daughter cell will exhibit the same signaling state as its parent in a given oscillation. Errors for observed results are standard errors. Zhai and colleagues do not give error rates for their calculations, so these are estimates. Errors for all simulated results are standard deviations. Consistency fractions are based on data from Larkin et. al (2018), with errors estimated as for a binomially distributed observation. For the signaling fraction and pairwise recurrences, these are across 20 runs. The rest are across five. Table S1 is an extended version of this table with data from inner cells and the total population, additional measures, and a description of the standard error estimation.

|  | Observed | Simulated |
| --- | --- | --- |
| Signaling Fraction | $0.43 \pm 0.02$ | $0.43 \pm 0.012$ |
| Signaler Recurrence | $0.60 \pm 0.1$ | $0.58 \pm 0.023$ |
| Non-signaler Recurrence | $0.78 \pm 0.1$ | $0.69 \pm 0.023$ |
| Consistent Signaling Fraction | $0.38 \pm 0.03$ | $0.37 \pm 0.005$ |
| Consistent Non-signaling Fraction | $0.50 \pm 0.03$ | $0.52 \pm 0.008$ |
| Inconsistent Fraction | $0.12 \pm 0.02$ | $0.11 \pm 0.004$ |

In our simulations, we also examined consistency across many waves of signaling and across an entire signaling oscillation, not just looking at a snapshot of signaling during the peak. Figure 4 shows cellular signaling consistency across 30 oscillations, with 5 replications. We found that 50% of cells consistently signaled (*>* 90% of the time), 44% consistently did not signal (*<* 10% of the time), and 6% were inconsistent, with a smaller mode at 50% participation among cells that signaled inconsistently. Note that this adds up to more than the mean of 43% signalers observed at oscillation peaks. This is due to the fact that more than 43% of cells signal each oscillation, but some signal before and some after each peak.

**Figure 4:**
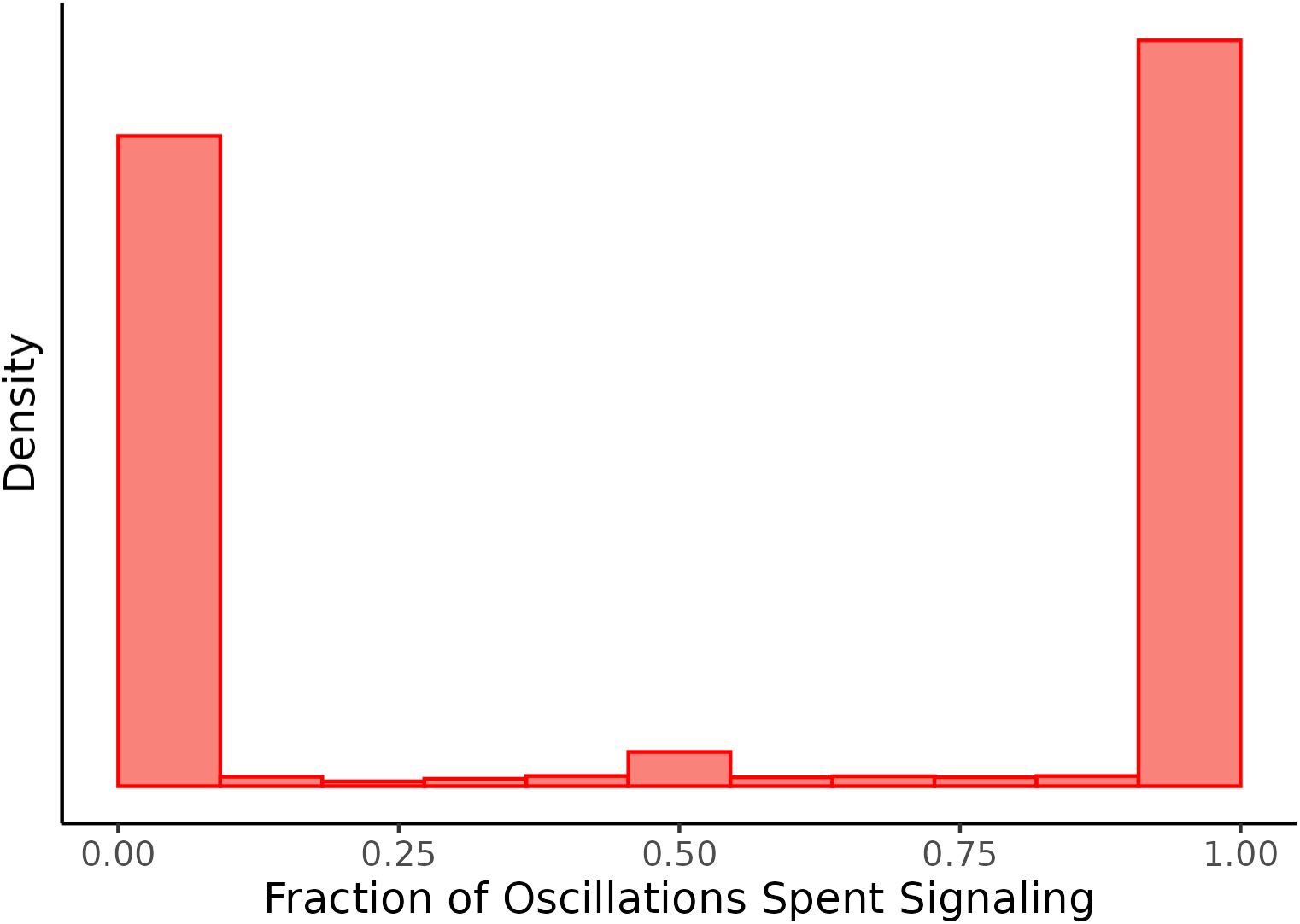
Histogram of the average number of signaling peaks during which cells signaled. Approximately 250,000 cells were tracked across 30 oscillations, and each bar in the histogram represents the number of cells that signaled in a proportion of signaling waves in the corresponding range. Peaks at one and zero indicate that most cells were consistent in signaling or not signaling (respectively).

Finally, Zhai et al. (2019) found that signaling behavior appears heritable—the daughter cells of cells that participate in signaling are more likely to participate in signaling themselves. In our model, the signaling thresholds of individual cells are noisily inherited, and this inheritance aligns with the observations of Zhai and colleagues. For example, with our selected values for signaling threshold inheritance, approximately 58% of daughter cells of signaling cells signal themselves, and approximately 69% of daughter cells of cells that do not participate in signaling also do not participate, close to the observations of Zhai et al. (Table 1). Further exploration of the effect of cell-level threshold on signaling appears in Supplement S2 and Figure S5. To maintain comparability to the findings of Zhai and colleagues, we measured concordance of signaling status for each mother-daughter pair during a peak of signaling (though different measures are given in Table S1).

#### Responses to media perturbations

*B. subtilis* biofilm oscillation experiments have taken place within a strictly controlled environment, where glutamate, as the only nitrogen source in the media, acts as a limiting nutrient. Liu et al. (2015) showed that, in such an environment, oscillations can decrease or stop in response to an increase in basal glutamate (the level of glutamate in the media surrounding the biofilm). Martinez-Corral et al. (2018) further found that oscillations would begin at a smaller biofilm size if basal glutamate were reduced, and showed that depolarization during biofilm growth can cause a wave of signaling. Figure 5 shows the results of simulations intended to replicate these findings in our model. By increasing basal glutamate, we weakened oscillations (Figure 5A). By drastically increasing potassium to depolarize the biofilm, we caused an initial peak of signaling (Figure 5B), and by lowering basal glutamate, we triggered early oscillations (Figure 5C).

**Figure 5:**
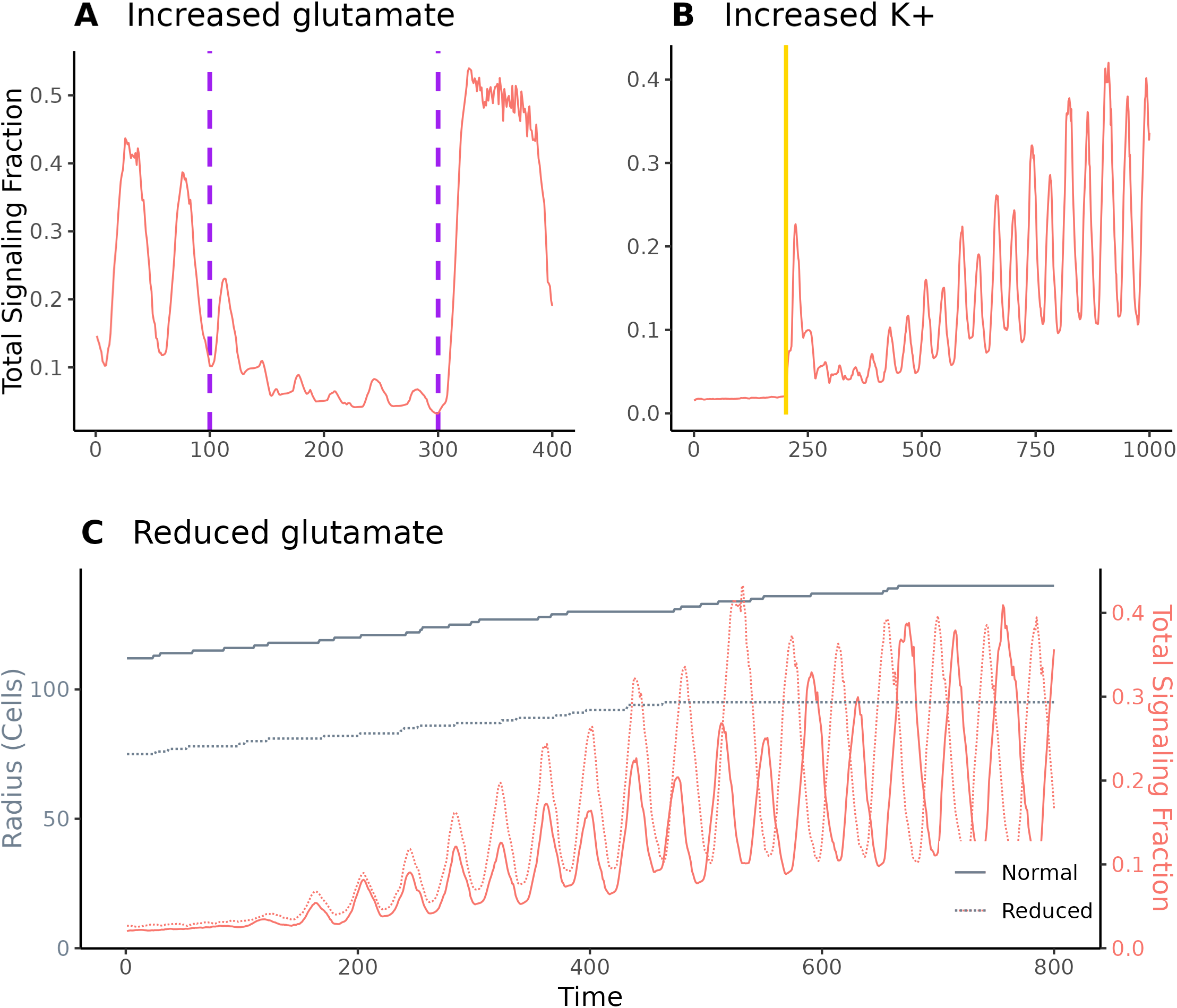
Effects of environmental conditions on signaling. (A) Increasing basal glutamate from 30 mM to 35 mM from ticks 100 to 300 in a biofilm that has been stably oscillating caused a depression in oscillation magnitude. (B) Depolarizing a growing biofilm by increasing basal potassium from 8 to 300 mM for five ticks (indicated by the gold band) caused a wave of signaling. This mimicked the methodology from Martinez Corral et al. (2018). (C) By growing a biofilm in a reduced-glutamate environment (G_m_ = 20 mM) we caused oscillations to begin at a much smaller population size. The radius for this biofilm levels off earlier because the oscillations will collapse if the biofilm grows to full size (Figure S2). Note that signaling rates in this figure are for the entire biofilm, not just outer cells, and are therefore sometimes higher than those reported elsewhere.

### Applications and predictions

#### Oscillation synchronization between adjacent biofilms

In addition to reproducing previously observed experimental results, our model can make predictions that motivate new experiments. Liu et al. (2017) found that two biofilms that are adjacent to each other will shift their oscillations to synchronize, but they did not identify a mechanism for this synchronization. Two molecules whose external concentrations are likely affected by depolarization waves are glutamate and potassium. To test whether our model could replicate synchronization and explore its explanation, we imposed external oscillations of both glutamate and potassium within our simulations. Our model parameters include basal levels of glutamate and potassium, so we simulated the effect of signaling in an adjacent biofilm by oscillating basal glutamate and basal potassium separately (Figure 6). Glutamate oscillations do lead the biofilm to synchronize, but only if the magnitude of glutamate oscillation is substantially greater than we would expect to be caused by a neighboring biofilm (Figure S6). In contrast, signaling oscillations change rapidly to be synchronized if basal potassium is oscillated even at relatively low magnitude. Our model therefore replicates the synchronizing behavior observed by Liu et al. (2017) and predicts that it is more strongly driven by neighboring biofilms’ effects on potassium than those on glutamate.

**Figure 6:**
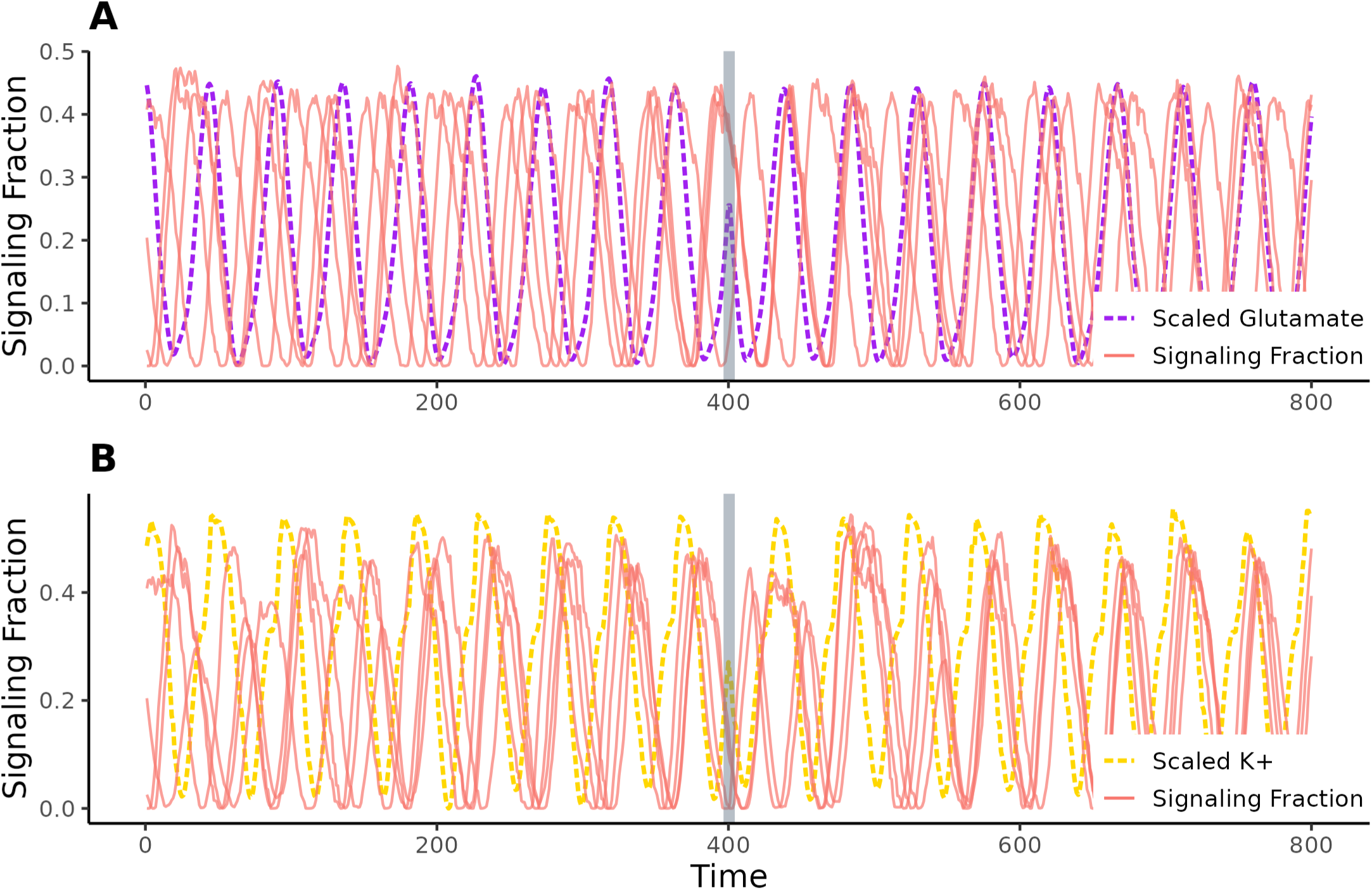
A comparison to determine whether the synchronization observed between adjacent biofilms is affected by (A) glutamate or (B) potassium ions. We oscillate basal glutamate (violet) by (−0.07, 0.1) mM and basal potassium (gold) by (−0.07, 0.06) mM following the trajectories of glutamate among exterior cells and external potassium respectively, taken from one of our simulations. After 400 ticks we accelerated the basal oscillation by a quarter period. Each solid red line indicates a different simulation. Glutamate oscillations do not appear to have a strong effect. However, when potassium is changed, the biofilm’s oscillations rapidly shift in response and remain closely synchronized across replicates.

#### Threshold effects

In our model, the propensity of a cell to signal is determined by its stress threshold. If a cell’s internal glutamate falls below its stress threshold, then the cell will signal. The results described above were simulated using thresholds distributed over a truncated normal distribution, with a mode on the parental-cell threshold, lower bound of 0, upper bound of 3, and *ff* of 1. To explore the effects of this distribution, we tested the signaling patterns and internal glutamate of biofilms across a variety of threshold bounds. We found that the distribution bounds must fall within a certain range in order for signaling to remain stable (Figures 7G, S7). If the maximum bound is too low, then signaling occurs, but only at very low levels (Figure 7A and D). There are never enough signalers to starve the exterior and trigger a wave of signaling, so only the interior cells signal. If the minimum bound is too high, then signaling collapses (Figure 7C and F). Too many cells signal simultaneously, and signaling is uncoordinated. All cells become stressed enough to signal and at any given time half or more are signaling. Between these regimes, the biofilm exhibits stable oscillations (Figure 7B and E).

**Figure 7:**
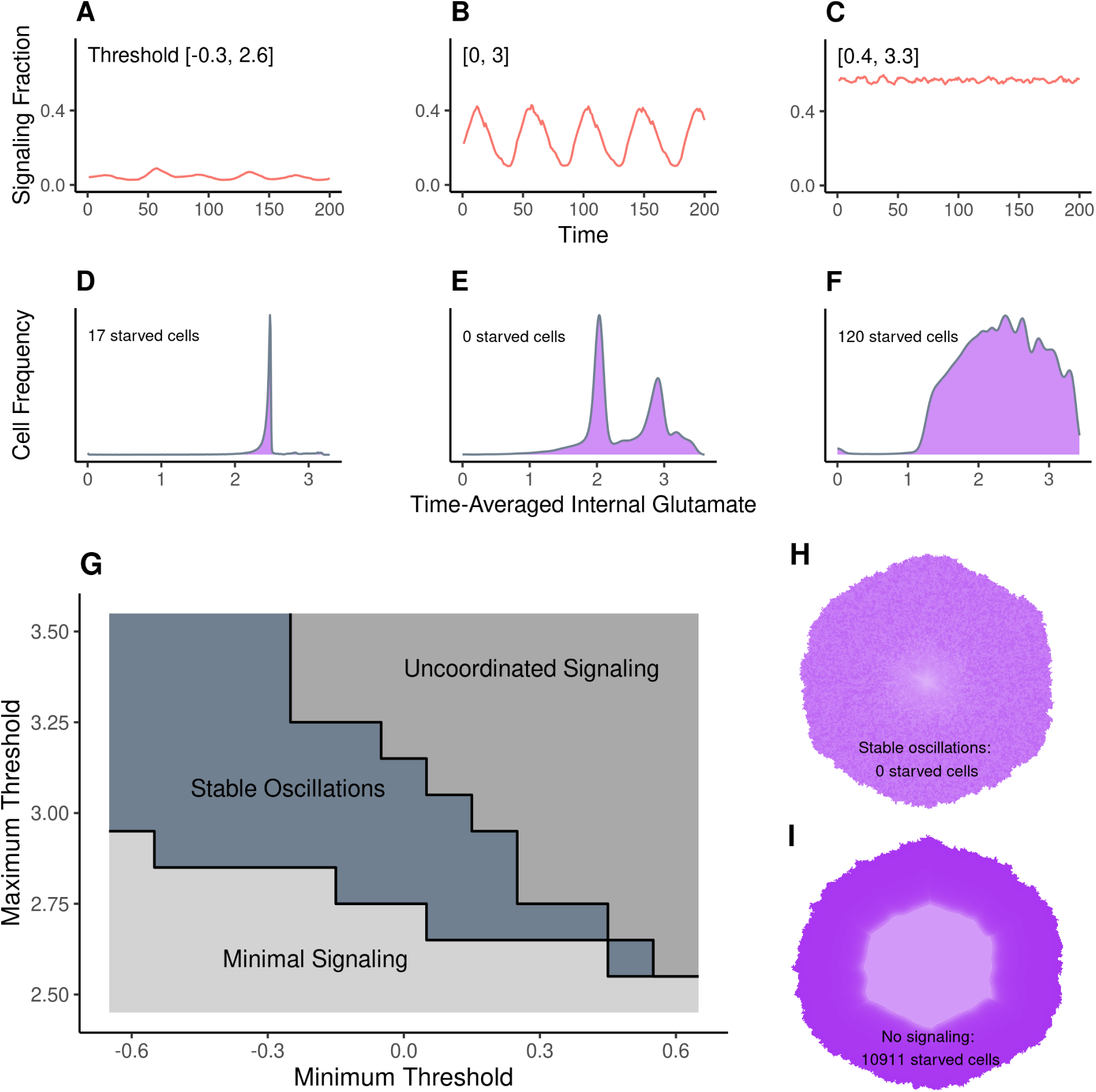
The effects of signaling threshold range on oscillations patterns and glutamate distribution. (A) shows the fraction of signalers over 200 ticks for a biofilm with low thresholds [−0.3, 2.6]. (Cells with stress thresholds ≤ 0 never signal; more negative values of the lower bound lead to more cells that never signal.) (D) displays the corresponding internal glutamate levels averaged across time for all cells in the biofilm. Seventeen cells starved—had less than 10^−5^ mM internal glutamate on average after the end of biofilm growth. (B) and (E) display the same for a range of [0, 3], and (C) and (F) for [0.4, 3.3]. (G) shows a phase diagram of the region of maximum and minimum signaling thresholds in which we observe stable oscillations. The region of stable oscillations produces oscillations with a range of more than 20% between the lowest level of signalers and the highest (eg. (B)). Minimal signaling indicates a low average level of signaling (as seen in (A)), and the region of uncoordinated signaling produces results like in (C). The trajectories for the simulations used to produce this phase plot are in Figure S7. (H) is the time-averaged internal glutamate for the biofilm in (B), dark purple indicating higher internal glutamate. (I) is the same, except for a biofilm with no signaling, leading to the interior 10,911 cells starving. Versions of (H) for the other two boundary conditions can be found in Figure S8.

It has been proposed that potassium signaling promotes an even distribution of glutamate across the biofilm, plausibly improving the survival rate of interior cells [25, 34]. We tested this idea by tracking the distribution of glutamate across cells in simulations that either did or did not include signaling behavior. By comparing the mean internal glutamate of cells across oscillations, we can see the effect of signaling. Without any signaling, exterior cells obtained substantial glutamate, but interior cells did not, with more than 10,000 (approximately 20% of all cells) reaching zero glutamate (Figure 7I). However, in simulated biofilms that signal, glutamate is much more evenly distributed across the biofilm, with zero cells having no glutamate (Figure 7H). Any amount of signaling produced substantially fewer starving cells (Figures 7A-C and S8), but only stable oscillations resulted in no starved cells. This suggests that potassium signaling does promote even distribution of glutamate by slowing growth and allowing glutamate to diffuse to interior cells, potentially increasing the stability of the biofilm during periods of high metabolic stress.

## Discussion

We introduced a computational model of metabolic signaling in *B. subtilis* biofilms. Previous models of this behavior have either been small in scope, only able to examine local behaviors of cells and omitting nutrients, or large in scope but unable to study heterogeneity in cell-level behavior [50, 11]. We have developed a model that bridges this gap, allowing the examination of the effect of cell-level behaviors on broader signaling patterns and the concentration of nutrients across the biofilm. We were able to replicate both individual-cell and biofilm-scale observations from previous work and new experiments, including oscillation and growth patterns, signaling in interior and exterior cells, and synchronization between neighboring biofilms. We also found support for the hypothesis that signaling results in a more even distribution of glutamate, which may extend the lifespan of a biofilm during periods of stress.

Previous models of *B. subtilis* signaling have adopted various assumptions about the effects of signaling on individual cells and the biofilm. On one hand, the models of Prindle and colleagues (2015), Martinez-Corral and colleagues (2018, 2019), and Ford and colleagues (2021) encoded assumptions that imply that signaling will increase glutamate uptake for the signaling cell both by directly increasing the cell’s ability to absorb glutamate, and suppressing glutamate absorption for neighboring cells.

On the other hand, the models in Larkin et al. (2018) and Zhai et al. (2019) prioritized the observation that hyperpolarized cells experience slower growth [24], although more recent work has suggested that the slow growth of hyperpolarized cells may be an artifact of ThT staining itself inhibiting growth [14]. Larkin and colleagues hypothesized signaling to be costly to the individual cell but beneficial to the biofilm as a whole, as it promotes a more even distribution of glutamate. Further, they noticed that the fraction of cells that signal in a given wave was close to the minimum number of cells necessary for the signaling wave to propagate across the exterior of the biofilm as predicted by percolation theory [41] (where signalers are randomly distributed among non-signalers and a signal is propagated by direct contact between two signaling cells). They interpreted this observation as being consistent with the idea that signaling cells act altruistically, sacrificing their own growth to promote the integrity of the biofilm.

In our model, we adopt assumptions similar to those of Prindle et al. (2015) and Martinez-Corral et al. (2019) that lead to signaling typically increasing the glutamate uptake of the signaling cell. At the same time, we replicate the heterogeneity in signaling behavior, the fraction of signaling cells, and the individual-level consistency of signaling across waves emphasized by Larkin et al. (2018) and Zhai et al. (2019). Thus, the individual-cell-level signaling patterns observed by the latter studies—and particularly a fraction of signaling cells near the percolation-theory threshold for signal transmission—can be attained without an explicit trade-off between individual-level growth and group-level glutamate distribution. However, like Larkin et al. (2018) and Zhai et al. (2019), our results are consistent with the idea that cell-level heterogeneity is important. In our model, a particular amount of variation in propensity to signal is necessary to achieve synchronized oscillations. In the presence of such variation, the cells with the highest propensity to signal hyperpolarize first. Once enough cells participate, a wave of signaling occurs, relieving glutamate stress and suppressing further signaling. Under this hypothesis, the participating fraction of cells may be near the level predicted by percolation theory because once that level is reached, stress is relieved and further signaling is not required.

The observation that a requisite level of variation in signaling propensity is necessary to produce coordinated waves of signaling in our model raises further questions. What could be the source of variation in signaling propensity, and how could this variation be maintained? *In vitro* biofilms observed to participate in signaling are typically clonal, so variation in signaling behavior is unlikely to be genetic in well-studied cases. Yet signaling behavior is observed to be heritable, in the sense that daughter cells are more likely to participate in signaling waves if their mother cell signals. One speculative possibility is that the regulatory network controlling potassium channel expression [27] results in multi-generational epigenetic inheritance of signaling [45, 32]. Whatever the source of the variation, on the basis of current observations, if the apparent individual-level cost of signaling is in fact an artifact of ThT staining [14], cells with a proclivity to signal might be expected to increase in frequency within the biofilm, taking up more glutamate than their neighbors, dividing more quickly, and potentially transmitting (non-genetically) their elevated propensity to signal to their offspring. Depending on how propensity to signal is realized and transmitted, such a process could lead to a decline of variation in propensity to signal, or at least to a decline of heritable variation, if continued long enough and if there are no forces generating new heritable mutation (analogous to mutation). (Our model contains such a force, as random deviations from a parent cell’s signaling threshold are partially inherited by offspring.) In our model, if too many cells signal, oscillations cease to be coordinated, and the distribution of internal glutamate—while much more even than in the complete absence of signaling—leaves some cells at the interior of the biofilm starved of glutamate. Thus, our model raises a possibility that is almost the reverse of the one raised by Larkin et al. (2018) and Zhai et al. (2019)—if signaling improves glutamate uptake for the signaling cell and reduces glutamate uptake for its neighbors, we might think of the cells that do not signal, rather than the ones that do, as acting altruistically, giving up their access to glutamate so that interior cells are not starved. There remain other possibilities—there may in fact be a cost of signaling to the individual, the increase in glutamate uptake from signaling may be dependent on the signaling state of a cell’s neighbors, or any of a number of others. In our current implementation, reproduction is not dependent on internal glutamate, so we do not explore such questions, but they are important for future theoretical and experimental work.

Another area of future study involves extending our model to predict how other processes are altered by emergent electrochemical signaling. For example, the expression of some genes has been proposed to be regulated by ion-responsive kinases [12]. By coupling cellular potassium flux to gene expression in our model, we could predict patterns of gene expression heterogeneity that would arise due to signaling. In addition, other cell phenotypes are regulated by nutrient conditions, notably matrix production and sporulation [26]. By modeling the response of genetic circuits that control the differentiation into these phenotypes [5], we could predict how the altered distribution of nutrients in signaling biofilms in turn alters the distribution of matrix producers and spores [46, 40, 6]. Our model may prove valuable to understanding the feedback between cellular phenomena and emergent nutrient conditions within biofilms, a topic of recent interest [17].

Overall, our work shows that combining agent-based and diffusion-based models can account for the emergence of community-level properties from interactions of individual cells. Doing so allows us to study the effect of signaling behavior on the biofilm as a whole, and on individual cells, taking into account heterogeneity among cells. That so many of the collective and cell-level signatures of *B. subtilis* biofilm signaling can be observed in a simple model hints at a relatively simple set of principles governing *in vitro* signaling behavior.

## Methods

### Model development

Our model is a network agent-based model, where cells are simulated as individual “agents,” each with their own set of rules for interacting with each other and their environment. Cells are placed on a network, where each cell is on a node and can interact with its neighbors. In the context of biofilms, neighbors are adjacent cells. During each unit of time (a “tick,” representing 1.2 minutes in this model) every cell performs actions according to their governing equations, and the environment is updated. We model the biofilm as hexagonal, matching observations by Larkin et al. (2018) that cells in these biofilms have a modal value of 6 immediate neighbors.

To determine which interactions to include and how cells should behave, we followed the model from Martinez-Corral et al. (2019). Their model is an ODE system describing a one-dimensional cross-section of a *B. subtilis* biofilm. We simplify their equations to be tractable for an agent-based model, leaving us with 4 equations (S1, S3-S5) that describe potassium uptake and release, glutamate uptake and consumption, membrane potential, and the interactions between potassium, glutamate, and membrane potential.

#### Initialization and growth

To initialize the model, we “grow” the biofilm, drawing each layer from the previous one. We begin by making a hexagon of 7 cells (6 outer and one center cell). These have signaling thresholds (the level of internal glutamate they can drop to before they will signal) randomly drawn from a uniform between 0 and 3. We then grow the biofilm to a radius of 50 cells while all external variables remain static: we ignore diffusion, metabolism, and signaling during this period. Each tick we randomly select one-fortieth of the cells on the perimeter of the biofilm network, with replacement, to reproduce. Each daughter cell (*j*) is a clone of its parent (*k*), except that its signaling threshold is drawn from a truncated normal with bounds of 0 and 3 in most of the work reported here, and with *ff* of 1 and *—* equal to the parent’s signaling threshold. The cell is placed in one of the empty nodes adjacent to the parent, with probability proportional to the number of neighboring cells each empty node has.

Once this initial phase of growth is complete, we begin to simulate potassium and glutamate behavior. Each tick, we update potassium via equation S4, simulating absorption, signaling, and diffusion. Simultaneously, we update glutamate via equations S1 and S3, simulating metabolism and absorption and using the algorithm described in Supplement S3.1 to approximate diffusion. We calculate the change in membrane potentials for each cell based on the results from the potassium calculations (equation S5). We continue growth at a rate of one-fortieth of the perimeter per tick until the network occupies 75% of the maximum size of ∼68,000 cells.

### Model validation

We validated our model by replicating previous experiments by other researchers. As a control, we ran the model 20 times under default conditions (using the parameters given in Table S2). Each run recorded a variety of data, with 5 of the runs recording individual signaling and glutamate data for every cell during each tick. These runs were used to gather summary statistics including signaling rate, recurrence rates and growth trajectories.

#### Perturbations

To test the effect of increased glutamate, basal glutamate was increased to 35 mM from 30 mM for 200 iterations in a biofilm that had already been growing for 2600 iterations. To test potassium shock, we increased basal potassium from 8 mM to 300 mM for 5 ticks in a growing biofilm, beginning at 750 ticks. We also simulated a biofilm with basal glutamate at 20 mM, limiting its growth to a radius of approximately 90 cells. The results from these perturbations are shown in Figure 5.

#### Oscillation synchronization

To explore the effect of a neighboring biofilm signaling in proximity to our simulation, we oscillated basal glutamate and potassium. To replicate the magnitude of change a signaling biofilm would cause in the surrounding media, we used the trajectory of external potassium and that of internal glutamate among exterior cells from a normal run of our model. These oscillated around their means by (−0.35, 0.32) mM and (−0.36, 0.48) mM respectively. We then scaled these by 0.2 to represent the effect of distance, for a final oscillation of (−0.07, 0.06) mM for potassium and (−0.07, 0.1) for glutamate. We oscillated each for 400 ticks, then skipped the oscillating molecule forward by a quarter period and simulated for another 400 ticks. These results are given in Figure 6. We also replicated these with more extreme scaling. Glutamate was oscillated by 200% scaling (−0.72, 0.96) and potassium by 5% (−0.018, 0.016). These results are reported in Figure S6.

### Experiments

Biofilms experiments were performed in a microfluidic device (CellASIC ONIX2 B04-F plate, Millipore Sigma, Burlington, MA, USA) as described in previous work [34, 20]. Cells (*Bacillus subtilis* strain NCIB3610, Bacillus Genetic Stock Center) were streaked on LB agar plates, incubated overnight at 37^◦^C, grown in liquid LB medium, resuspended in liquid msgg medium for additional growth, and loaded into the microfluidic plate. The composition of msgg was 5 mM potassium phosphate (pH 7.0), 100 mM MOPS (pH 7.0), 2 mM MgCl_2_, 700 *—*M CaCl_2_, 50 *—*M MnCl_2_, 100 *—*M FeCl_3_, 1 *—*M ZnCl_2_, 2 *—*M thiamine HCl, 0.5% (v/v) glycerol and 0.125% (w/v) monosodium glutamate. After cell loading into the microfluidic plate, biofilms were grown under flow at 30^◦^C and Thioflavin-T (ThT) was added to the media for imaging cellular membrane potential after 12 hours of growth [34]. Biofilms were imaged in phase contrast and fluorescence with a 4X, 0.13 NA objective on an Olympus IX-83 microscope (Evident Scientific, Waltham, MA, USA).

Time traces of ThT were extracted from time-lapse movies using a machine learning-based segmentation approach implemented in Python, which applies a Random Forest classifier, provided by the Scikit-learn library, trained on manually segmented biofilm images to perform segmentation using the ThT fluorescence channel. In the ThT traces of Figure 3, we subtracted slow accumulation of ThT *post hoc* to make oscillation traces stationary.

Pairwise signaling consistency calculations given in Tables 1 and S1 were calculated by tracing the signaling states of approximately 300 cells across a 2 hour period that included two oscillations.

## Code availability

Code used to generate the simulations and figures that appear in this manuscript is available at https://github.com/Muldero/AgentBasedBsubtilis.

## Acknowledgments

We thank C. Bergstrom, T. Kessinger, M. Lachmann, C.B. Ogbunu, J. Van Cleve, and members of the Edge, Mooney, Pennell, and Larkin labs for helpful comments on this study. Funding was provided by NIH grant R35GM137758 to MDE and NIH grant R35GM142584, NSF grant 2027108, and a CASI award from the Burroughs Wellcome Fund to JWL.

