## Supplementary Information for "An Agent-Based Model of Metabolic Signaling Oscillations in *Bacillus subtilis* Biofilms"

### Supplementary text for ”An Agent-Based Model of Metabolic Signaling Oscillations in *Bacillus subtilis* Biofilms”

### S1 Definitions

#### S1.1 Signaling

The phenomenon we refer to as “signaling” occurs via membrane polarization. Larkin et al. (2018) observed that cell polarization during a “wave” of signaling is bimodal, with cells that have recently released potassium having high polarization and other cells having lower polarization. We define signaling status similarly to Larkin and colleagues. When we examine membrane potential directly, signaling is not clearly bimodal, but when transformed by Equation S2 ($\mathscr{F}(V) = \frac{1}{1+e^{V-V_0}}$) it becomes bimodal (Figure S4, transformation taken from Martinez-Corral et al. (2019 equation S3)). We then define signaling cells as those having $\mathscr{F}(V)$ greater than the median of the two modes, and non-signaling cells as those having $\mathscr{F}(V)$ less than the median of the two modes.

#### S1.2 Interior and exterior cells

Both our simulations and *in vitro* observations show that the interior and exterior of the biofilm behave differently from each other (Figure 3). We track behavior for these two groups of cells separately in addition to tracking biofilm-wide indicators. The boundary between inner and outer cells is not determined visually, nor by signaling behavior. Instead, we observed that cell behavior changes qualitatively at approximately the depth to which glutamate diffuses if there is no signaling. To determine the depth from the exterior of the biofilm to the boundary, we thus set cell membrane potentials to be the default of -156 mV, internal potassium to its default of 300 mM, and external potassium to the basal level of 8 mM. We then apply our algorithm for determining glutamate diffusion and set all cells that receive any glutamate as exterior. Those at a depth to which glutamate does not diffuse are defined as interior cells. This method provides a slightly conservative estimate, as the boundary generated is somewhat closer to the exterior of the biofilm than the qualitative boundary between inner and outer cells. This approach is appropriate to ensure consistency for our summary statistics of outer cells with those from prior studies that focused solely on the biofilm's exterior.

### S2 Changing the heritability of the signaling threshold

Changing the standard deviation of the truncated normal from which signaling thresholds were drawn changed the heritability of signaling behavior. The lower the standard deviation, the more heritable the signaling threshold. If $\sigma$ was reduced too much, then the distribution of thresholds across the biofilm became uneven and strongly dependent on the initial draws of signaling thresholds at the foundation of the simulated biofilm. In this regime, signaling usually occured either at a very low rate or in an uncoordinated way. If $\sigma$ was raised to 2, then we still observed normal oscillations of approximately the same magnitude as if $\sigma$ equals 1, but the signaling recurrence rates decreased appreciably, as might be expected. The daughters of signaling cells signaled 49 $\pm$ 1% of the time, as opposed to 58 $\pm$ 1% when $\sigma = 1$ and 60% in the *in vitro* observations of Zhai et al. (2019). For non-signaling cells, daughter cells' frequency of non-signaling behavior dropped from 0.69 $\pm$ 0.01 to 0.59 $\pm$ 0.01, as opposed to 0.78 *in vitro*.

### S3 Model description

Our model was built on the equations in the following subsections, adapted from Martinez-Corral et al. (2019). Martinez-Corral and colleagues modeled glutamate and potassium diffusion, absorption, release, and metabolism. They also modeled glutamate membrane transporter density, ammonium (which affects the rate of decay of intracellular glutamate) and GDH enzyme (which affects the production of ammonium). The price of using an agent-based model is that simulations are slower and more computationally expensive, so we applied some simplifications. We modeled glutamate, potassium, and membrane potential, but we did not model transporters, ammonium, or GDH. Instead we set those parameters to the default values from Martinez-Corral et al. (2019). The parameter values for these equations can be found in Table S2. Further, for the factors we did explicitly model, we closely followed the approach in Martinez-Corral et al. (2019).

#### S3.1 Glutamate

Internal glutamate ($G_i$) is the amount of glutamate within each cell. Each tick, we calculated glutamate absorption (affected by the uptake constant $\alpha_g$, the membrane potential $V$, and the amount of external glutamate $G_e$). Internal glutamate is consumed at a constant rate $\delta_g$. The higher the external glutamate, the higher the rate of glutamate uptake—the rate of uptake depends on $k_g$, the concentration of extracellular glutamate at half-maximal uptake rate. We assumed that cells do not release glutamate, and so the rate of change of internal glutamate is the rate of uptake minus consumption,

$$\Delta G_i = \alpha_g \mathscr{F}(V) \frac{G_e}{k_g + G_e} - \delta_g G_i \tag{S1}$$

where

$$\mathscr{F}(V) = \frac{1}{1 + e^{V - V_0}}. \tag{S2}$$

Here $\mathscr{F}(V)$ represents the logistic effect of membrane potential on glutamate uptake. If a cell is polarized, then there is minimal hindrance to uptake, but once it becomes depolarized, glutamate uptake rapidly goes to zero.

External glutamate for each cell (the amount of glutamate outside of the cell, used to determine glutamate uptake) is affected by glutamate uptake, which moves external glutamate to the cell's interior, and by diffusion. To make our model computationally tractable, we modeled glutamate diffusion via a simplified one-dimensional approximation. We assumed that glutamate diffused exclusively from outside the biofilm through to the middle, with no diffusion perpendicular to this direction (i.e. no tangential diffusion around the 2D biofilm). For perimeter cells, we set $G_e$ to the basal concentration of glutamate. We then calculated glutamate absorption for perimeter cells and used the remaining glutamate as the initial $G_e$ for the next layer. This process was repeated layer by layer, averaging the leftover glutamate from the outer layers. Interior layers of the biofilm contain fewer cells than exterior layers. Thus, on moving in one layer, we scaled the amount of external glutamate by the ratio of the number of cells in the adjacent interior layer to the number of cells in the current layer. In this way, glutamate approximately diffuses to the biofilm center each tick, but is allocated amongst cells radially. Labeling the amount of glutamate diffusing inward to a given cell from the adjacent exterior layer as $D_g$, the change in external glutamate is given by

$$\Delta G_e = D_g - \alpha_g \mathscr{F}(V) \frac{G_e}{k_g + G_e}, \tag{S3}$$

where the second term is the same glutamate absorption term that appears in equation S1.

#### S3.2 Potassium

In our model, internal potassium concentration is governed by potassium uptake and by potassium release that occurs during signaling. Potassium uptake is proportional (via a constant $\alpha_k$) to the product of internal glutamate, external potassium, and the difference between a homeostatic set point for potassium ($K_{i0}$) and the cell's current internal potassium ($K_i$). (Note that we labeled this process "uptake," but it can reverse in principle if $K_i > K_{i0}$.)

When a cell signals (encoded below by the indicator variable $\mathcal{I}_T$; when a cell is signaling, $\mathcal{I}_T = 0$, otherwise $\mathcal{I}_T = 1$), it releases a substantial amount of potassium, with the amount depending on the levels of interior and exterior potassium. (Generally there is much more potassium inside the cell than outside, and cells with more internal potassium, relative to external potassium, release more potassium when signaling.) The amount of potassium released also depends on the the membrane polarization of the cell, $V$ (which is typically negative). Combining uptake with outflow during signaling, the change in internal potassium is given by

$$\Delta K_i = \alpha_k G_i K_e (K_{i0} - K_i) - F g_K \left( V - V_{K0} \ln \frac{K_e}{K_i} \right) \mathcal{I}_{\{T\}}, \tag{S4}$$

where $F$ is the membrane capacitance, $g_K$ is the potassium channel conductance, $V_{K0}$ is the Nernst potential prefactor, and $\mathcal{I}_T$ is an indicator variable that equals 1 if the cell is currently signaling and 0 otherwise.

Like glutamate, external potassium is determined by the uptake and release of potassium from a cell, plus diffusion. Unlike glutamate, potassium is a small ion and diffuses rapidly with respect to the coarse discretization of time we employed [4, 3]. Further, potassium has a much more even distribution across the biofilm than glutamate does. Thus, to simulate diffusion, we computed per-cell external glutamate levels by accounting for uptake and signaling via equation S4, and then averaged external potassium across all cells in the biofilm every tick.

#### S3.3 Membrane potential

When a cell is not signaling, the membrane potential approaches a set point $V_{L0} = -156$ mV, but can become more positive (less polarized) if external potassium is close to or greater than basal potassium in the media ($K_m$). During a signaling event, membrane potential becomes much more negative (polarized) in proportion with the amount of potassium released. Following Martinez-Corral et al. (2019), we modeled changes in membrane potential as

$$\Delta V = -g_L \left( V - \left( V_{L0} + d_L \frac{K_e - K_m}{1 - e^{(K_m - K_e)/\sigma}} \right) \right) - g_K \left( V - V_{K0} \ln \frac{K_e}{K_i} \right) I_{\{T\}}. \quad \text{(S5)}$$

This expression is a discrete version of equation S18 of Martinez-Corral et al. (2019), with the signaling indicator variable $I_T$ replacing a variable indicating propensity to signal. $g_L$ is the leak conductance and $d_L$ is the leak slope coefficient.

#### S3.4 Parameters

Most of the model parameters we used were taken from Martinez-Corral et al. (2019). For basal nutrient concentrations, the values were calibrated to equal those in their *in vitro* experiments. The rest of the parameters used in equations S1:S5 were borrowed from Martinez-Corral et al. (2019) as well but do not have as strong a basis in observations or theory.

##### S3.4.1 Scaling

Our model is in units of ticks, while Martinez-Corral et al. (2019) used hours (as have other ODE-based approaches). We aimed to set ticks to last 1.2 minutes and based the growth rate and parameter values off of this. Thus all parameters in units of hours were scaled by a factor of 50. However, in our model, tick size does not change the rate at which glutamate diffuses because we assumed that it diffuses approximately to the center of the biofilm each tick. This assumption was used to make the model computationally tractable. But it actually takes around 4.1 minutes for glutamate to diffuse to the center of the biofilm ([4] and our own calculations), inducing a discrepancy with our other assumptions. To account for this, we adjusted the parameters dictating potassium uptake $\alpha_k$ and membrane capacitance $F$ (by factors of 1.5 and $3.1^{-1}$ respectively). The fact that glutamate diffusion is slower than we assumed may account for the faster-than-expected oscillations displayed by our model, which have a period of approximately an hour, as opposed to approximately two hours on average *in vitro*. Increasing the tick size to match the 4.1 minutes it takes glutamate to diffuse seemed to be too coarse for the model to tolerate, and changes in many of the variables between ticks become large, leading to unstable behavior. While developing the present model, we pursued another, more computationally intensive approach that involved simulating glutamate and potassium diffusion explicitly, giving some behaviors that matched the current results and oscillation periods closer to *in vitro* observations (results not shown). We opted for the current model for its simplicity and lower computational burden.

### S4 Supplemental figures

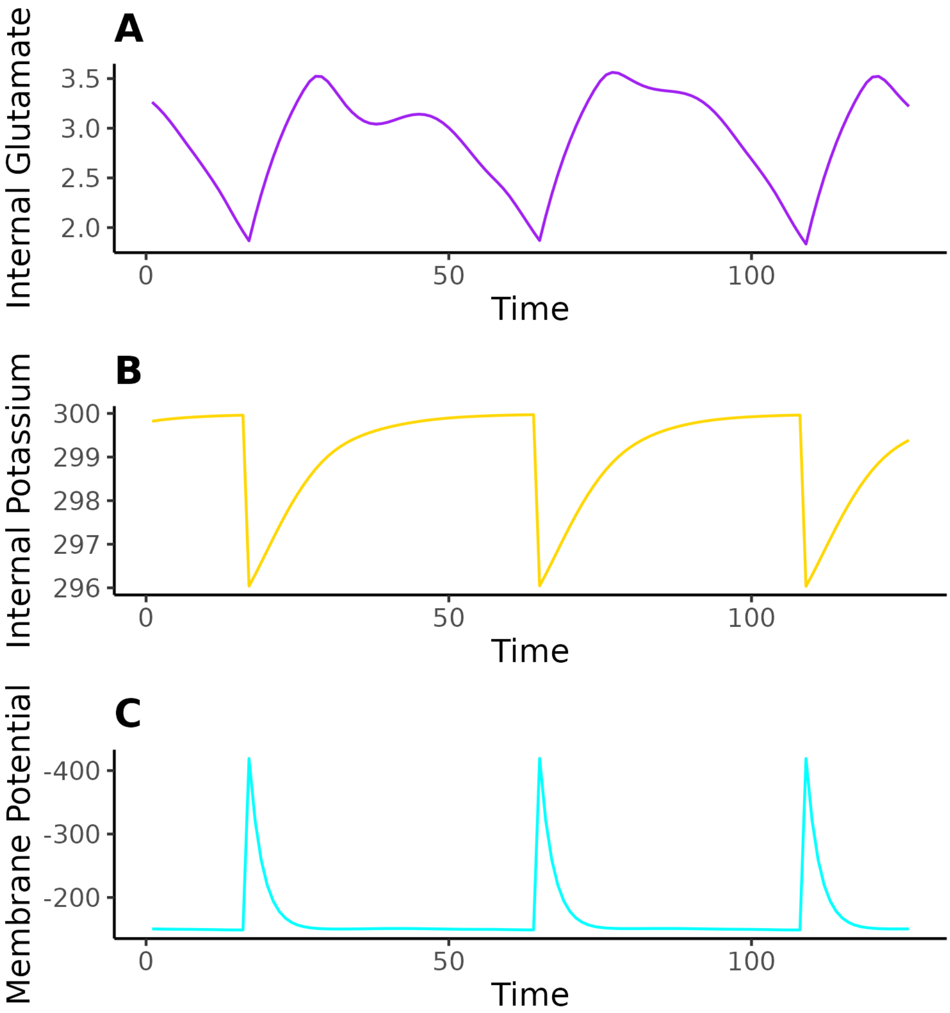


*Figure S1: Time-series data for a signaling cell in the periphery of the biofilm. (A) Internal glutamate is initially consumed faster than glutamate uptake can replenish it. Once internal glutamate drops below a threshold value, signaling occurs, and the cell absorbs more glutamate. (B) When signaling occurs, a cell releases potassium, causing internal potassium levels to drop. After that, potassium slowly returns to its set point. (C) The release of internal potassium during signaling causes membrane potential to spike temporarily, facilitating glutamate uptake. Oscillations take about 45 ticks each. Potassium and glutamate are in mM, membrane potential is in mV.*

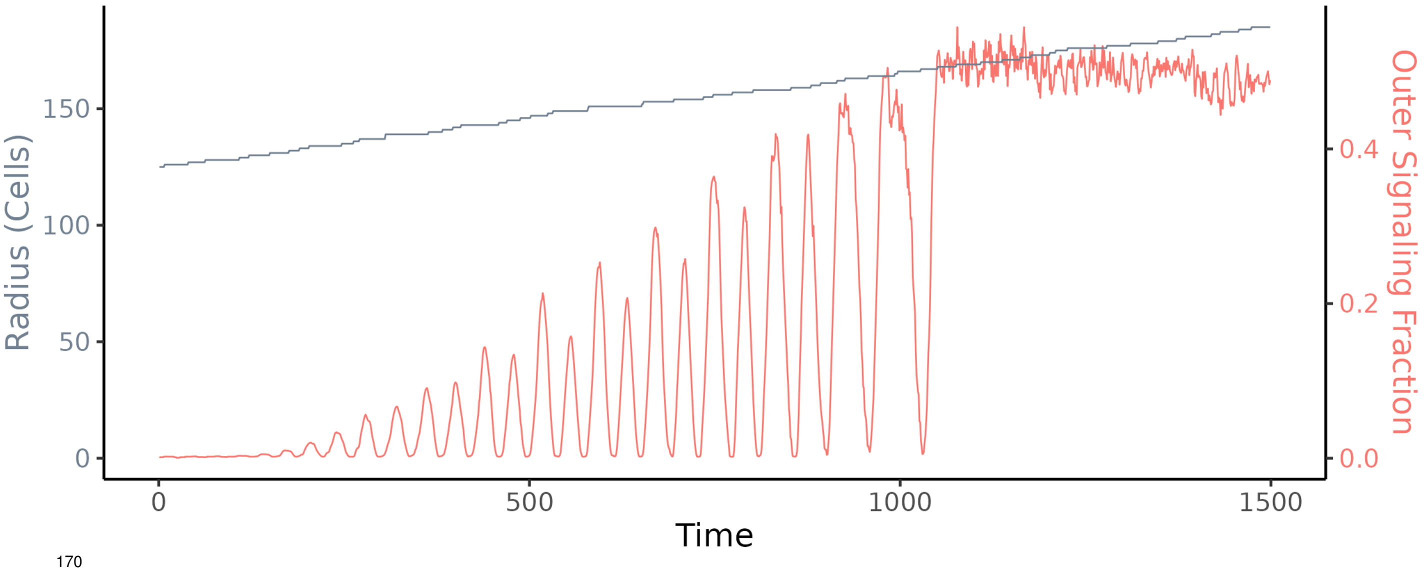


*Figure S2: This is an alternative version of Figure 3 in which we set the maximum radius to be approximately 230 cells instead of 145. The fraction of signalers increases with the radius of the biofilm until the biofilm becomes too large and signaling oscillations collapse (around tick 1100). Cells continue to signal, but signaling is randomly distributed across the biofilm, with most cells taking part.*

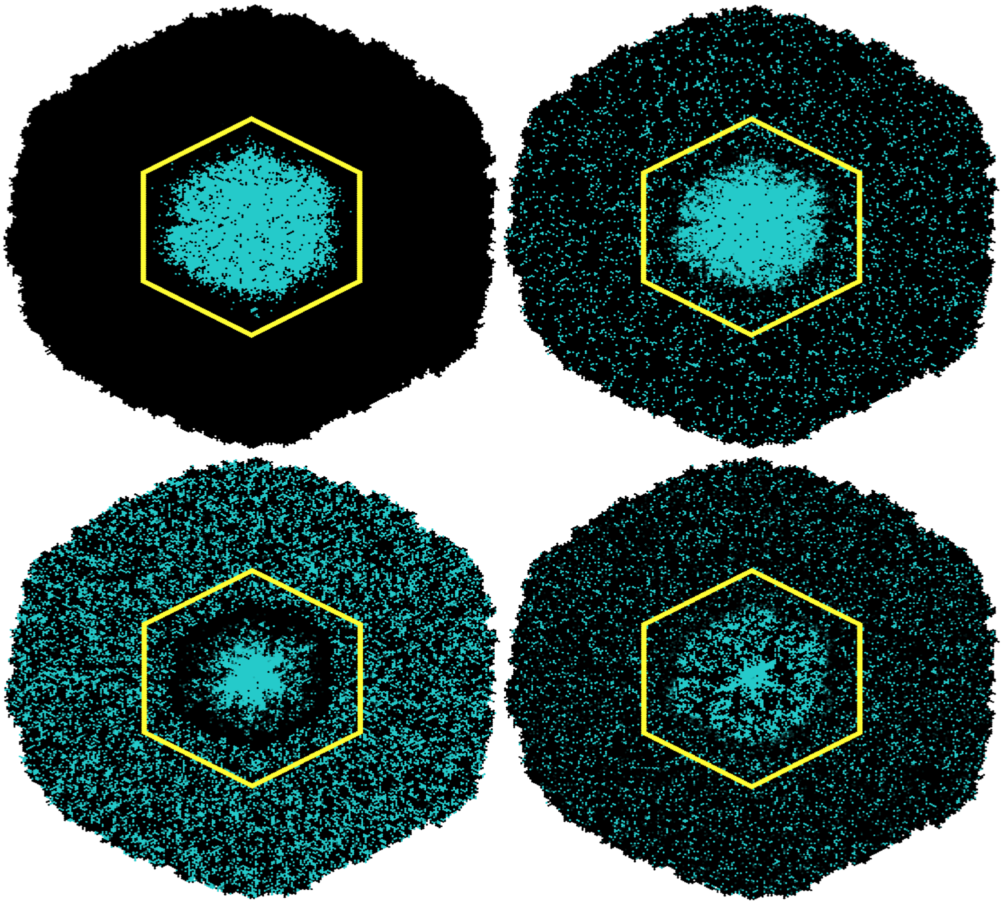

*Figure S3: An illustration of the difference in behavior between the interior and exterior of the biofilm across different time points in an oscillation, and our definition of the boundary (the yellow hexagon). In defining the boundary, we chose to err toward misclassifying exterior cells as interior. This is because we wish to compare the behavior of exterior cells in our model to previous observations, and thus want no interior cells included in the exterior. Cyan indicates hyper-polarized cells.*

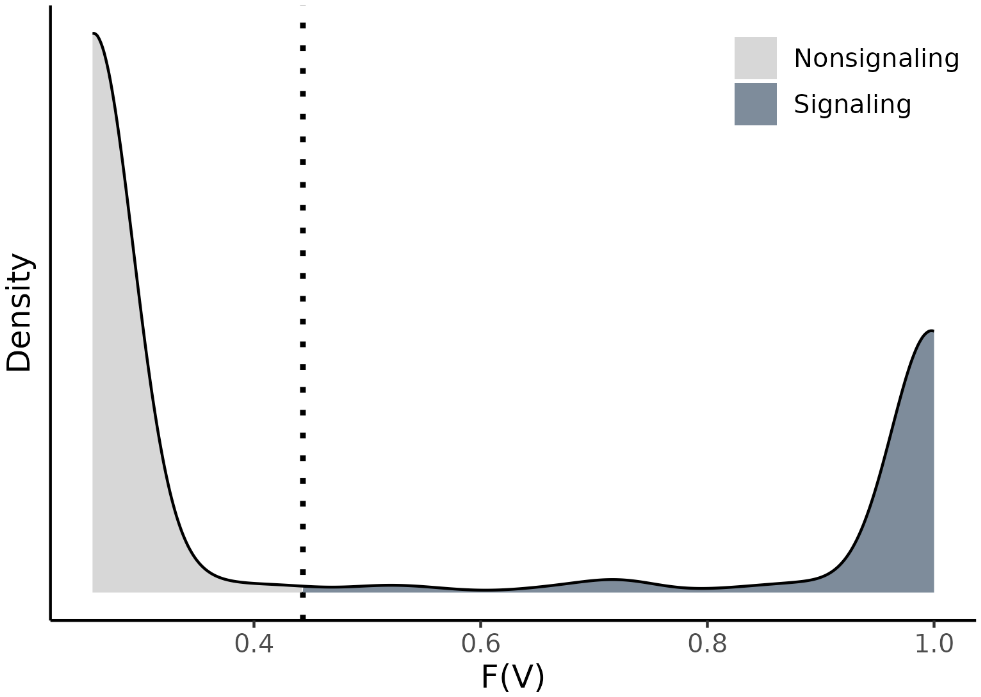


*Figure S4: Distribution of logistically transformed cell membrane potentials during a signaling peak. The dotted line indicates the point one quarter of the way between the lower mode and the upper mode. Cells with a transformed membrane potential greater than this (dark gray) are defined as signalers. Those with one lower (light gray) are non-signalers.*

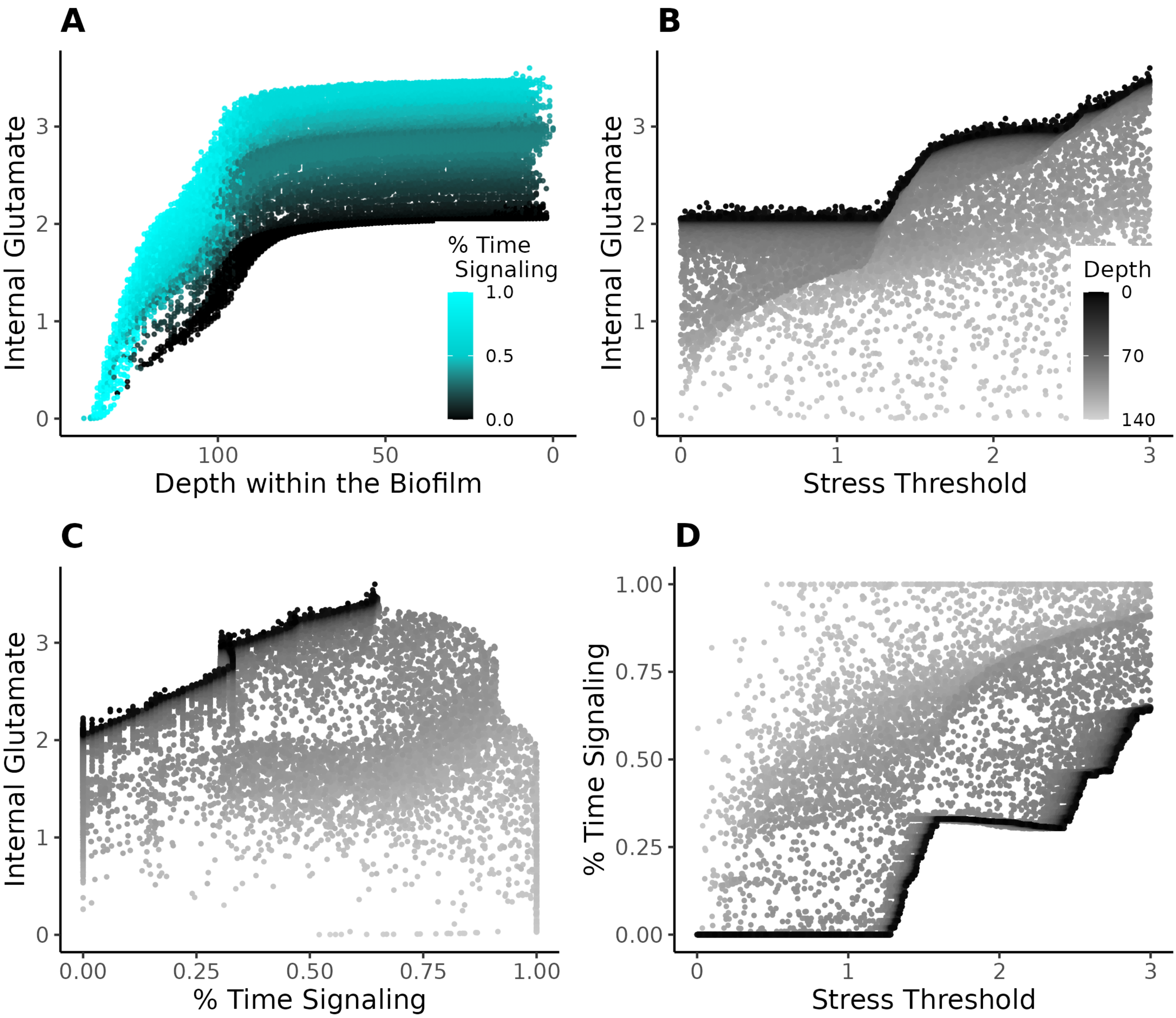


*Figure S5: A variety of visualizations comparing internal glutamate, cell-level signaling thresholds, percentage of time spent signaling (by cell and across all ticks, not just during signaling peaks), and depth within the biofilm (0 indicating a cell on the edge of the biofilm). The dark line of cells in (B-D) indicates cells on the exterior of the biofilm, and the scattered lighter cells are interior cells.*

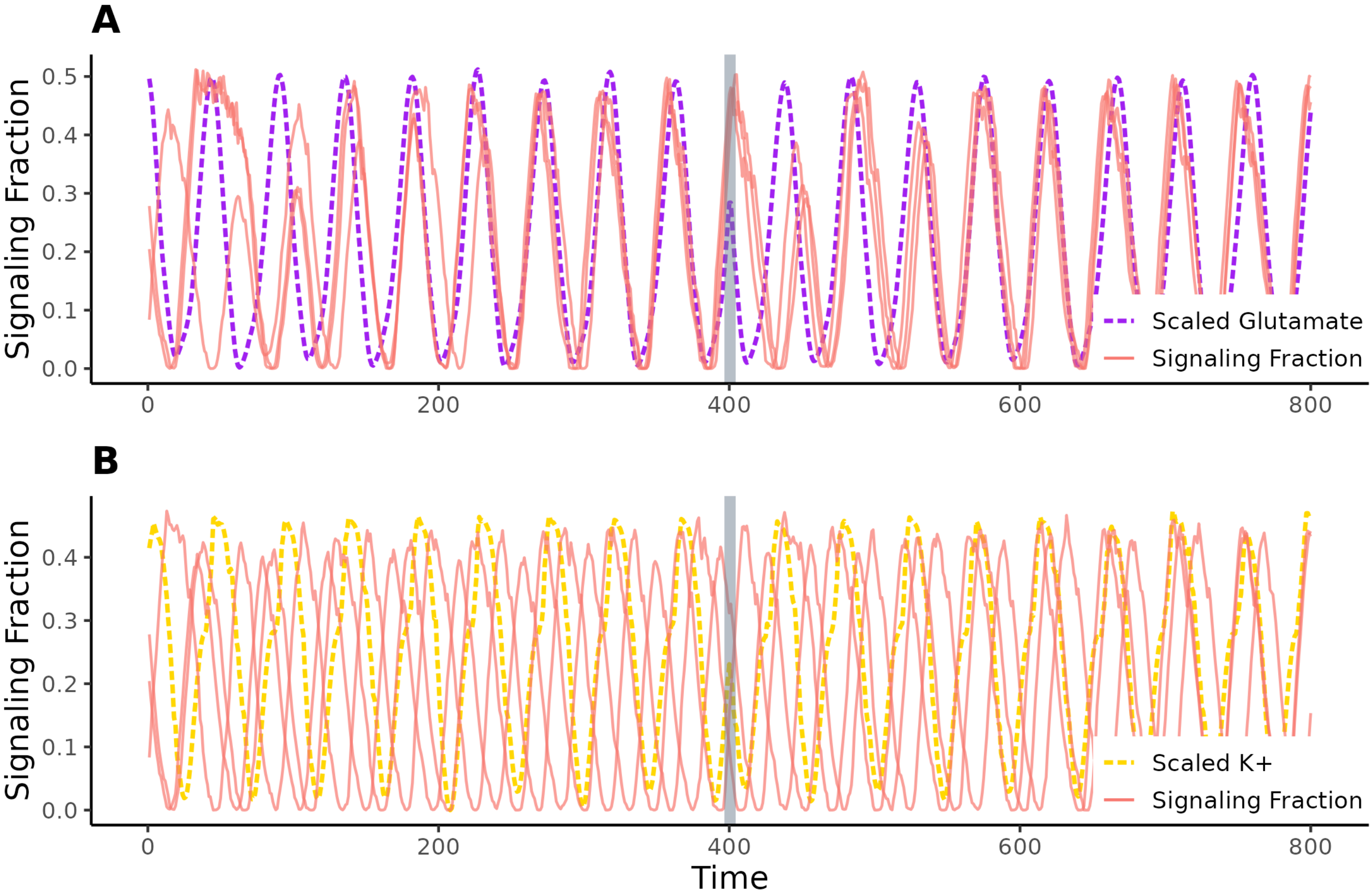


*Figure S6: An alternative version of Figure 6. In that figure we demonstrated that a small perturbation of 20% the observed variation in external potassium to mimic a nearby biofilm; this was enough to cause oscillation synchronization. An oscillation of 100% the internal glutamate variation was not enough to cause strong synchronization. Here we replicated that experiment but with 5% potassium and 200% glutamate. With these more extreme values we observe opposite effects, with glutamate causing rapid synchronization but potassium having a much weaker effect on synchronization.*

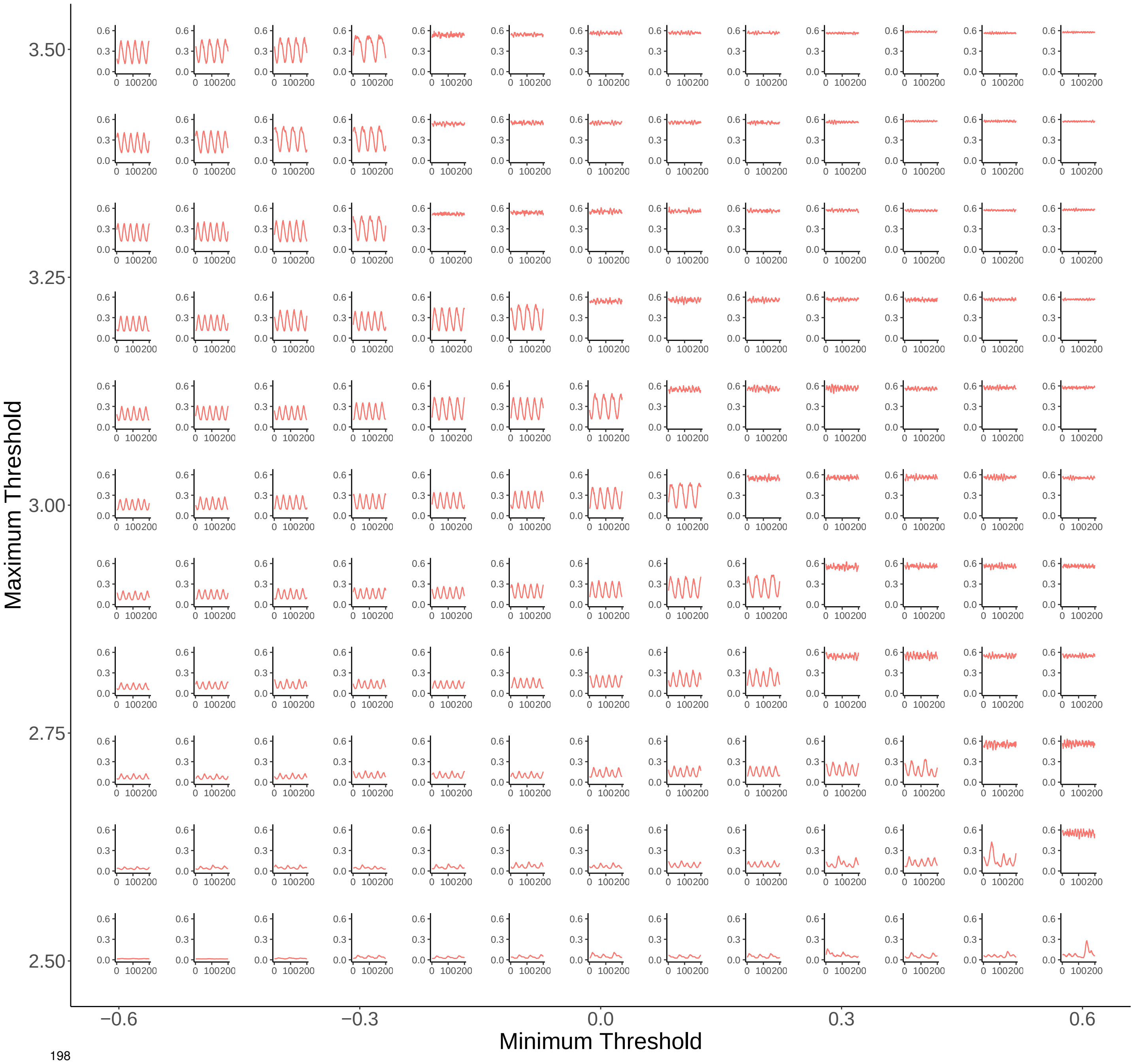


*Figure S7: The behavior of signaling oscillations for simulations across various signaling threshold ranges. The threshold bounds for each simulation are [x, y], rounded to the nearest 0.1. For each sub-plot, the x axis is the time (200 ticks) and the y axis the fraction of signaling cells. These data were used to create the phase diagram in Figure 7.*

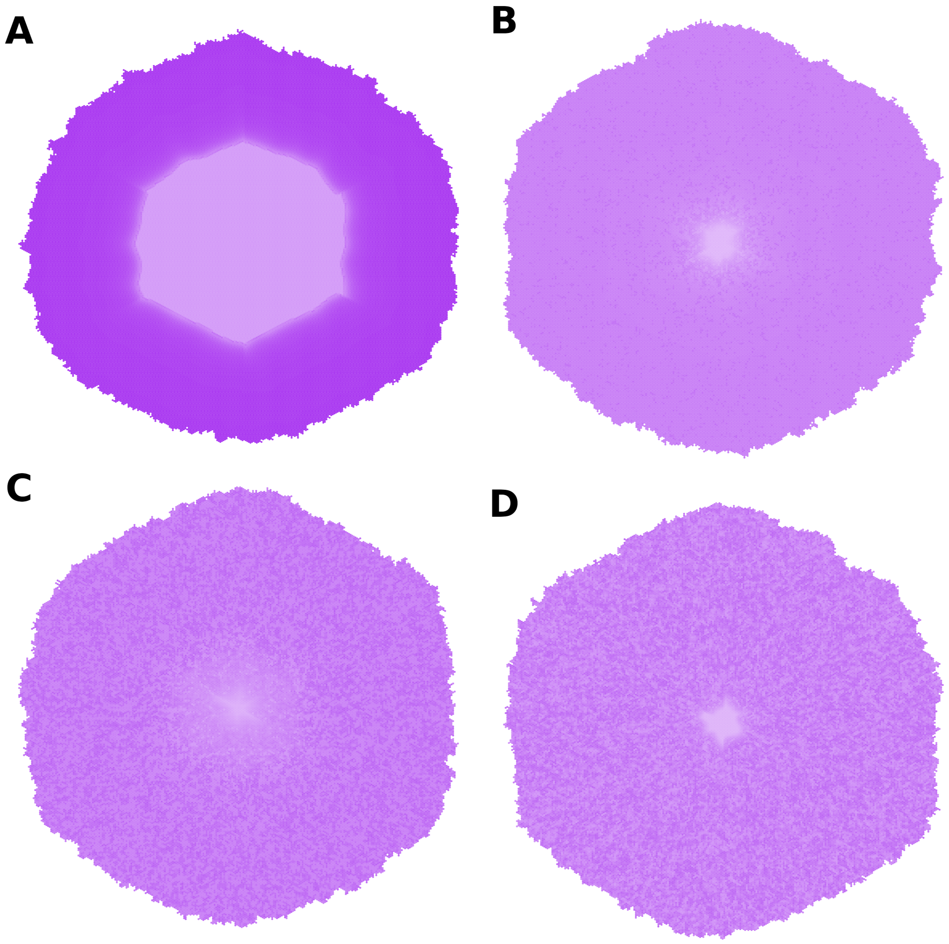


*Figure S8: An extension of Figure 7H and I. Here we show the mean internal glutamate over time for each cell in the biofilm. dark purple indicates high glutamate, light indicates low. (A) has no signaling. (B) has bounds of [$-0.3$, 2.6], producing minimal oscillations. (C) is the regime used in our main results—[0, 3]—which produces stable oscillations similar to those observed in vitro. And (D) has bounds of [0.4, 3.3], which produces a high level of uncoordinated signaling.*

### S5 Supplemental Videos

#### S5.1 *In vitro* time-lapse video

This is a time-lapse of microscope images of a *B. subtilis* biofilm exhibiting the oscillatory behavior *in vitro*. The biofilm has been stained with the fluorescent membrane potential reporter ThT; cyan indicates hyperpolarized cells. The scale bar is 100 $\mu m$.

*[See file "Video_S5.1.gif"]*

#### S5.2 Simulated time-lapse video

This is a time-lapse of our simulated model. Here too, cyan indicates hyperpolarization. This video loops after two oscillations.

*[See file "Video_S5.2.gif"]*

#### S6 Supplemental tables

| | Observed | Outer | Inner | Total |
|---|---|---|---|---|
| Signaling Fraction | 0.43 ± 0.02 | 0.43 ± 0.012 | 0.54 ± 0.022 | 0.41 ± 0.010 |
| Pairwise Signaler Recurrence | 0.60 ± 0.1[†] | 0.58 ± 0.023 | 0.84 ± 0.009 | 0.59 ± 0.020 |
| Pairwise Non-signaler Recurrence | 0.78 ± 0.08[†] | 0.69 ± 0.023 | 0.79 ± 0.017 | 0.71 ± 0.019 |
| Total Signaler Recurrence | NA | 0.51 ± 0.012 | 0.88 ± 0.014 | 0.75 ± 0.015 |
| Total Non-signaler Recurrence | NA | 0.71 ± 0.004 | 0.63 ± 0.019 | 0.67 ± 0.009 |
| Pairwise Consistent Signaling Fraction | 0.38 ± 0.03 | 0.37 ± 0.005 | 0.50 ± 0.005 | 0.36 ± 0.005 |
| Pairwise Consistent Non-signaling Fraction | 0.50 ± 0.03 | 0.52 ± 0.008 | 0.44 ± 0.006 | 0.53 ± 0.008 |
| Pairwise Inconsistent Fraction | 0.12 ± 0.02 | 0.11 ± 0.004 | 0.06 ± 0.001 | 0.11 ± 0.004 |
| Total Consistent Signaling Fraction | NA | 0.50 ± 0.002 | 0.68 ± 0.003 | 0.54 ± 0.002 |
| Total Consistent Non-signaling Fraction | NA | 0.44 ± 0.006 | 0.25 ± 0.003 | 0.40 ± 0.006 |
| Total Inconsistent Fraction | NA | 0.06 ± 0.005 | 0.06 ± 0.006 | 0.06 ± 0.004 |

*Table S1: An extended version of Table 1 in the text. This includes values for inner and total cells as well as outer cells. We also include consistency and inheritance estimates for the entire duration of signaling, not just pairwise, and with signaling measured across an entire oscillation, not only during the peak of signaling. Errors for all simulated results are standard deviations. For the signaling fraction and pairwise recurrences, these are across 20 runs. For the rest, they are across 5. Error for the observed signaling fraction is a standard error from Zhai et al. (2019). Errors for observed signaling consistency are standard errors estimated as for a binomially distrbuted observation (number of cells = 316).*

*[†]Standard error values were not reported in Zhai et al. (2019). We estimated them based on the given recurrence rates (0.6 and 0.78), fraction of signalers (43%), and number of observations (49 pairs of cells). This suggested 22 initial signalers, of whom 13 have signaling offspring, and 27 non-signalers, of whom 21 have non-signaling offspring. This produced the standard errors given above assuming binomially distributed counts.*

| Parameter | Description | Default Value | Units |
|---|---|---|---|
| $t_{pp}$ | Tick size | 50 | hours$^{-1}$ |
| $gpt$ | Generation per tick | 1/40 | generations/tick |
| $\alpha_g$ | Glutamate uptake constant | 24/50 | mM/($\mu$M t) |
| $k_g$ | Extracellular glutamate concentration at half-maximal uptake rate | 0.75 | mM |
| $\delta_g$ | Glutamate degradation constant | 4.8/50 | mM$^{-1}$ t$^{-1}$ |
| $\alpha_k$ | Potassium uptake constant | 0.18/50 | mM$^{-2}$ t$^{-1}$ |
| $K_{i0}$ | Homeostatic potassium setpoint | 300 | mM |
| $F$ | Membrane capacitance | 0.016 | mM/mV |
| $g_K$ | Potassium channel conductance | 70/50 | t$^{-1}$ |
| $V_{K0}$ | Nernst potential prefactor | 100 | mV |
| $g_L$ | Leak conductance | 18/50 | t$^{-1}$ |
| $V_{L0}$ | Basal leak potential | -156 | mV |
| $d_L$ | Leak slope coefficient | 4 | mV/mM |
| $\sigma$ | Leak threshold sharpness coefficient | 0.1 | mV |
| $V_0$ | Resting membrane potential | -150 | mV |
| $K_m$ | Basal potassium in media | 8 | mM |
| $G_m$ | Basal glutamate in media | 30 | mM |
| $l_t$ | Lower threshold bound | 0 | mM |
| $u_t$ | Upper threshold bound | 3 | mM |
| $\sigma_t$ | Threshold standard deviation | 1 | NA |

*Table S2: Default values and units for all parameters used in the model. Content, formatting, and descriptions closely follow those in Martinez-Corral et al. (2019). The parameter that was labeled $D_p$ in their paper has been changed to $\alpha_k$. Parameters divided by 50 were originally in units of hours but have been converted to be in units of ticks (t).*
