## Supplementary figures and images for "An Agent-Based Model of Metabolic Signaling Oscillations in *Bacillus subtilis* Biofilms"

### Video S5.1

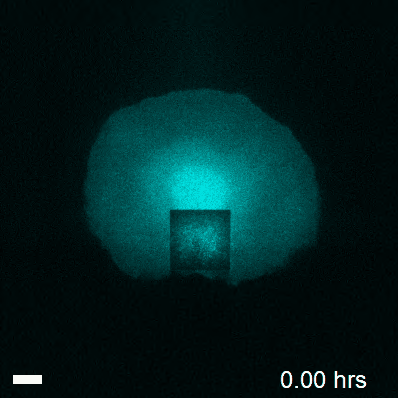

### Video S5.2

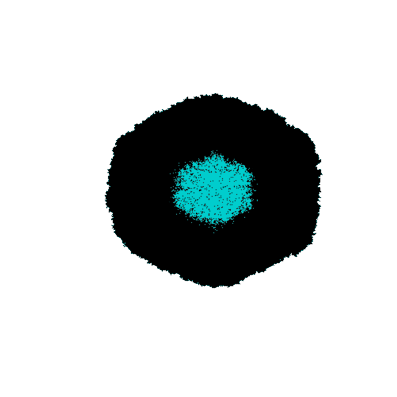
